# Multimodal classification of neurons in the lateral septum

**DOI:** 10.1101/2024.02.15.580381

**Authors:** Christopher M. Reid, Yuqi Ren, Rhiana Simon, Yajun Xie, Miguel Turrero Garcίa, Diana Tran, Steve Vu, Jonathan Liu, Manal A. Adam, Sarah M. Hanson, Angela O. Pisco, Corey C. Harwell

## Abstract

The lateral septum (LS) is a ventral forebrain nucleus that modulates complex social and affective behaviors. These behaviors emerge from heterogeneous neuronal populations whose molecular identity and developmental origins remain poorly defined. We profiled the transcriptional identity of mature LS neurons derived from two progenitor lineages distinguished by their embryonic origin and Nkx2.1 expression history, identifying 22 molecularly distinct subtypes. Nkx2.1-lineage neurons are enriched for select cell adhesion and communication molecules; however, subtypes from distinct developmental origins can converge onto similar molecular profiles when residing within the same LS subregion. The graded expression of genes related to synaptic signaling is a primary axis defining the taxonomy of LS neurons. Using transcriptional markers, we labeled non-overlapping neuronal populations and characterized their connectivity, morphology, and electrophysiology. Together, these findings define the extent of LS neuronal diversity and provide a framework for understanding how complex behaviors are regulated by the LS.

## INTRODUCTION

The ability to properly regulate affective and motivated behaviors is a fundamental requirement for social interaction and decision-making. These behaviors are orchestrated by a complex network of structures collectively known as the limbic system (Catani et al., 2013; Rajmohan and Mohandas, 2007; Sheehan et al., 2004; Sokolowski and Corbin, 2012). A complete understanding of how this system operates in diseased and healthy states requires a comprehensive knowledge of the cell types and circuitry of limbic structures. The lateral septum (LS) is a ventral forebrain region within the limbic system that plays a pivotal role in regulating complex affective, motivated, and social behaviors (Besnard and Leroy, 2022; Menon et al., 2022; Rizzi-Wise and Wang, 2021; Sheehan et al., 2004; Wirtshafter and Wilson, 2021). Compared to other limbic regions such as the cortex, hippocampus, and hypothalamus, our knowledge of the cell types present in the LS is limited (Cembrowski et al., 2016; Kim et al., 2020; Moffitt et al., 2018; Zeisel et al., 2015).

The LS is primarily composed of GABAergic projection neurons which have historically been classified based on their morphological and physiological properties, as well as their anatomical connections (Alonso and Frotscher, 1989; Gallagher et al., 1995a; Leranth and Frotscher, 1989; Risold and Swanson, 1997). These features have been used to subdivide the LS into three distinct nuclei: the lateral septum dorsal (LSd), the lateral septal intermediate (LSi), and the lateral septum ventral (LSv). Inputs coming from the hippocampus are topographically organized, with neurons located in dorsal areas projecting to the LSd and those in ventral regions projecting to the LSv. Similarly, fibers projecting from the LS to diencephalic and subpallial structures typically follow a lateral-to-medial trajectory, with the LSd projecting laterally and the LSv projecting medially (Raisman, 1966; Risold and Swanson, 1997; Swanson and Cowan, 1979).

Recently, groups of functionally distinct LS neurons have been characterized based on differences in the expression of neuropeptides and neurotransmitter receptors. For instance, somatostatin (*Sst)*-expressing neurons in the LSd have been implicated in the regulation of context-associated fear behaviors and are thought to be anxiolytic (An et al., 2022; Besnard et al., 2019; Li et al., 2022). In contrast, neurons expressing the corticotropin releasing hormone receptor 2 (*Crhr2*) in the LSi are activated by threatening stimuli and are anxiogenic (Anthony et al., 2014; Hashimoto et al., 2022). In the rostral LS, a group of neurotensin *(Nts)-*expressing neurons were shown to modulate social behaviors and gate hedonic feeding behaviors (Azevedo et al., 2020; Chen et al., 2022; Li et al., 2023). Collectively, these findings highlight the functional heterogeneity of LS neurons; however, a proper characterization of defined neuronal types requires a multimodal examination of their cell biological properties (Gouwens et al., 2020; Zeng, 2022).

Observations from well-studied areas of the brain such as the cortex have shown that the taxonomy of neurons within a region often corresponds to their developmental origin (Mayer et al., 2018; Tasic et al., 2016; Yao et al., 2021). The septal proliferative zone is characterized by the expression of ZIC family transcription factors (Gaston-Massuet et al., 2005; Inoue et al., 2007; Magno et al., 2017; Turrero García et al., 2021). Within this zone, progenitors in the caudal part of the embryonic septum—termed the septal eminence—can be differentiated from their rostral counterparts by the expression of the transcription factor *Nkx2.1* (Magno et al., 2017; Turrero García et al., 2023, 2021; Wei et al., 2012). Moreover, a loss of *Nkx2.1* in the septal eminence results in a severe depletion of medial septal and globus pallidus cholinergic neurons born from there (Magno et al., 2017). We previously described a subset of morphologically heterogeneous LS neurons originating from the septal eminence that are necessary for the proper execution of threat response behaviors (Turrero García et al., 2023, 2021). Taken together, these prior studies suggest that LS neurons with a developmental history of *Nkx2.1* expression may exhibit characteristics that distinguish them from those derived from the rostral embryonic septum.

In this study, we identified 22 transcriptionally distinct subtypes of LS neurons, which can largely be divided into two groups based on their developmental expression of *Nkx2.1*. We show that while neurons from the same lineage express common sets of cell adhesion and cell signalling molecules, ultimately developmental history does not fully predict molecular taxonomy. Additionally, we show that LS neuron subtypes are discretely organized into distinct domains along the dorsal-ventral and rostral-caudal axes of the septum. Furthermore, the transcriptional relationships between subtypes are defined by patterns of gene expression that correlate with the anatomy of the LS. Leveraging our transcriptomic data, we identified markers that can be used to characterize anatomical and physiological attributes of genetically defined non-overlapping subgroups of LS neurons. Collectively, this research provides a comprehensive examination of neuronal heterogeneity within the LS and lays the foundation for targeted functional studies.

## RESULTS

### Molecularly distinct lateral septal neurons exist within Nkx2.1+ and Nkx2.1− lineages

We reasoned that lateral septal neurons (LSNs) derived from Nkx2.1-expressing progenitors in the septal eminence may share cellular and molecular characteristics that correspond to their lineage origin (Sandberg et al., 2016; Turrero García et al., 2021; Wei et al., 2012). Neurons with a developmental history of *Nkx2.1* expression were labeled by crossing Nkx2.1-Cre mice with an Ai14 allele, a red fluorescent protein Cre reporter line. This marked a subset of LSNs (Nkx2.1+ LSNs) that were found in all anatomical regions of the LS and were positive for the ZIC transcription factors (**Figure 1A, S1A**). Using the neuronal marker NeuN, we quantified the total proportion of LSNs that were RFP-positive (22.7%), as well as their enrichment across the dorsal, intermediate, and ventral subregions of the LS (LSd: 11.7%, LSi: 29.2%, LSv: 29.7%) (**Figure 1B**). We also assessed the proportion of RFP-positive LSNs across four coronal sections along the anterior-posterior axis, finding them to be more concentrated in intermediate areas (Section 1: 15.4%, Section 2: 31.0%, Section 3: 27.3%, Section 4: 13.0%) (**Figure 1C**). The majority of Nkx2.1+ LSNs were positive for ZIC transcription factors; however, a small fraction (11.8%) lacked ZIC expression (**Figure S1B**), suggesting that these neurons may derive from Nkx2.1-expressing progenitors outside the embryonic septum, such as the medial ganglionic eminence or preoptic area (Azzarelli et al., 2015; Mayer et al., 2018; Turrero García and Harwell, 2017; Wei et al., 2012).

**Figure 1:**
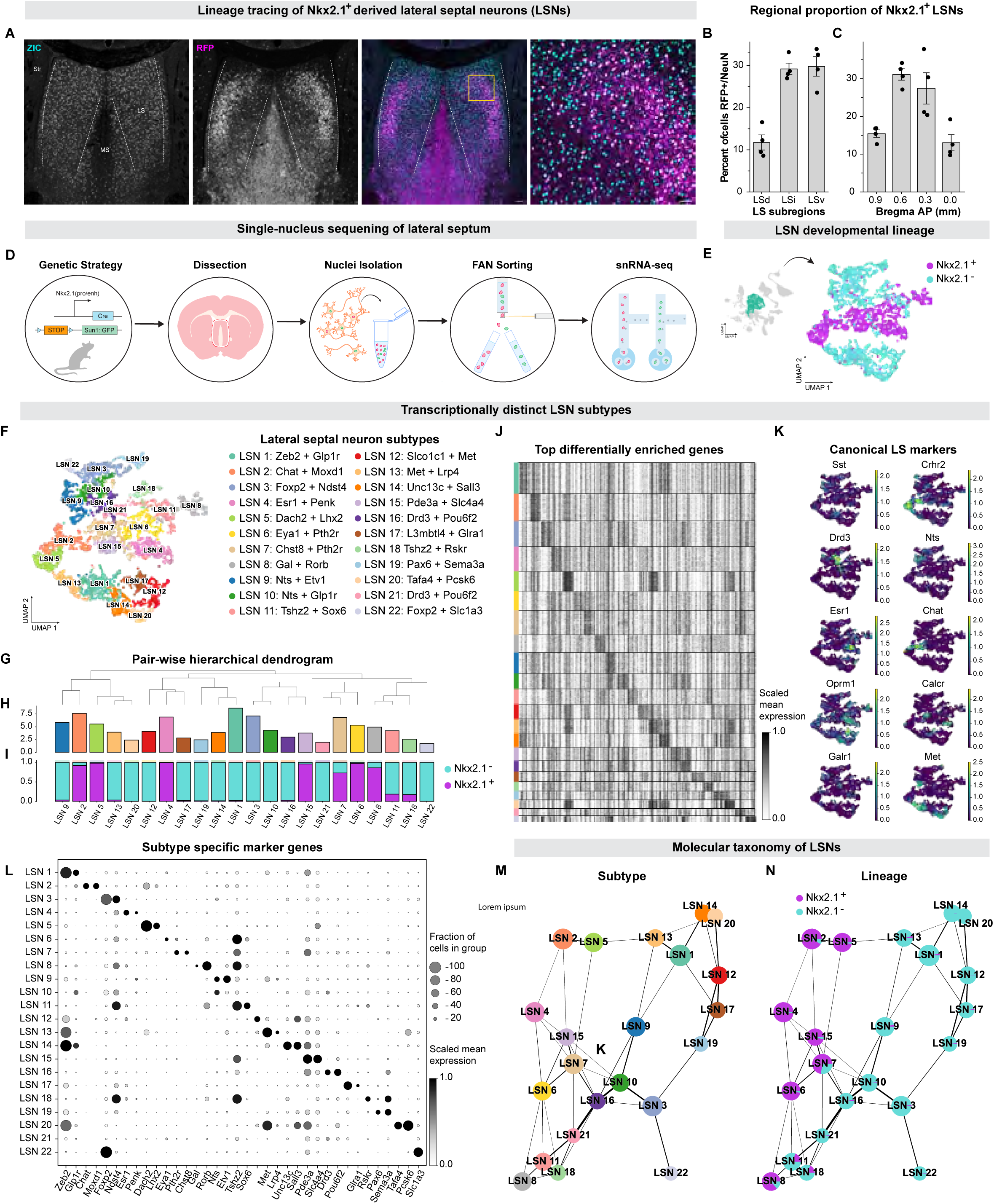
Transcriptional taxonomy of lateral septal neuron subtypes. **A.** Coronal section of the lateral septum immunofluorescently stained for RFP (magenta) and ZIC (cyan) in a P35 Nkx2.1-Cre animal carrying the Ai14 reporter allele. Scale bar: 100 μm **B.** Bar graphs showing the proportion of NeuN positive cells that are RFP positive (magenta) (N = 4). The left graph shows counts for the dorsal, intermediate, and ventral subregions of the LS. Right graph shows the counts for four regions along the anterior-posterior axis; from Bregma section 1 (S1): +0.9, section 2 (S2): +0.6, section 3 (S3): +0.3, section 4 (S4): 0.0. **C.** Schematic detailing our genetic strategy and single nucleus-sequencing approach. **D.** UMAP plot showing the GFP identity of 5,103 LSNs. **E.** UMAP plot of transcriptionally distinct LSN subtypes. **F.** Stacked bar graph showing the proportion GFP positive (magenta) or negative (cyan) cells in each LSN subtype. On top is a dendrogram based on hierarchical clustering analysis using the average total gene expression of each LSN subtype. **G.** Heatmap showing the top 10 differentially enriched genes for each LSN subtype. **H.** Bar plot showing the proportion of cells per subtype group. **I.** Dot plot showing the scaled average gene expression (scale bar) of the minimal markers necessary to define each LSN subtype. **J.** Feature plots showing the scaled expression (scale bar) of marker genes uniquely enriched in LSN subtypes. (**K-L).** PAGA based constellation plots showing which groups are similar to each other based on their molecular identify, annotated in by their lineage and subtype identities.

To determine if *Nkx2.1* expression history distinguishes LSNs, we developed an experimental strategy to profile the transcriptional identity of LSNs while preserving information about their developmental origins (**Figure 1D**). We dissected the septum of adult Nkx2.1-Cre mice carrying a conditional Sun1GFP allele for Cre-dependent labeling of the nuclear membrane (Mo et al., 2015). We then isolated the nuclei of septal cells and used fluorescent activated nuclei sorting (FANS) to separate GFP-positive and negative nuclei. Subsequently, we performed single nucleus RNA-seq (snRNA-seq) on each sorted population and obtained a total of 20,678 cells that expressed an average of 3,002 unique genes (Figure S2A). Using principal component analysis, we constructed a neighborhood graph which allowed us to visualize the data in two dimensions (**Figure S2B**). Our sequencing approach enabled us to distinguish between GFP-positive cells, indicating a history of *Nkx2.1* expression (Nkx2.1+) and GFP-negative cells that had never expressed *Nkx2.1* (Nkx2.1-) (**Figure S2C**).

We applied unsupervised Leiden clustering at several resolutions to understand how the groups of cells relate to each other (**Figure S2D**). 14 broad cell types were annotated using established marker genes that were indicative of their cell type, regional location, and developmental origin, including *Nkx2.1*, *Zic1*, and *Lhx8* (Mayer et al., 2018; Turrero García et al., 2021; Turrero García and Harwell, 2017; Yao et al., 2023; Lein et al., 2007) (**Figure S2E-G**). Our dissection methods captured cells from regions outside of the LS, such as medial septal neurons (MSNs), triangular septal neurons (TRNs), and striatal neurons (StrNs). Consistent with previous studies, we found that the majority of MSNs (93.6%) have a history of *Nkx2.1* expression (**Figure S2H and I**) (Magno et al., 2017; Wei et al., 2012).

We subclustered the 3,897 LSNs, which were primarily defined by their enriched expression of *Myo5b*, *Trpc4*, and *Prdm16* (Tian et al., 2014; Turrero García et al., 2021; Yao et al., 2023). UMAP analysis of their lineage identities revealed a clear segregation between Nkx2.1+ and Nkx2.1− LSNs (**Figure 1E**). To assess the heterogeneity of LSNs, we employed Leiden clustering at various resolutions (**Figure S3A**), aiding our annotation of 22 transcriptionally distinct subtypes (**Figure 1F**). To examine transcriptional similarity across groups, we hierarchically clustered the neuronal subtypes based on their average gene expression profiles and calculated their proportional representation, and fraction of Nkx2.1+ and Nkx2.1− LSNs within each group **(Figure 1G-I)**. This revealed that Nkx2.1+ and Nkx2.1− LSNs do not segregate into distinct clades, and that subtypes from non-overlapping lineages can share related molecular identities.

We then performed differential gene expression analysis to identify markers defining each subtype noting that many previously recognized markers, such as *Sst*, *Crhr2*, *Drd3*, and *Nts*, were selectively expressed by specific LSN subtypes (**Figure 1J and K)** (Anthony et al., 2014; Besnard and Leroy, 2022; Chen et al., 2022; Shin et al., 2018). Each subtype could be defined by a unique combination of selectively expressed genes (**Figure 1L, S3B**). Notably, we found that the graph-based embeddings of the LSN subtypes showed that Nkx2.1+ and Nkx2.1− divided into separate neighborhoods, suggesting broad programs exist within each lineage (**Figure 1M and N**).

### Nkx2.1−lineage neurons share expression of select cell adhesion and communication genes

To explore whether Nkx2.1+ and Nkx2.1− LSNs have distinguishing transcriptional programs, we conducted a pairwise differential gene expression analysis between the two groups, revealing 684 differentially expressed genes (157 enriched in Nkx2.1+ and 527 enriched in Nkx2.1− LSNs; **Figure 2A**). We found that the long non-coding RNA *Sfta3-ps* is highly enriched in Nkx2.1+ cells in a similar pattern to *Nkx2.1*, whereas the transcription factor gene *Meis2* is predominantly expressed in Nkx2.1− cells (**Figure 2B**). The mutually exclusive expression patterns of the two genes reflected the developmental origins of the LSNs born from either the rostral or caudal embryonic septum, dividing the subtypes into two groups: those that express *Meis2* and those expressing *Sfta3-ps* (**Figure 2C**).

**Figure 2:**
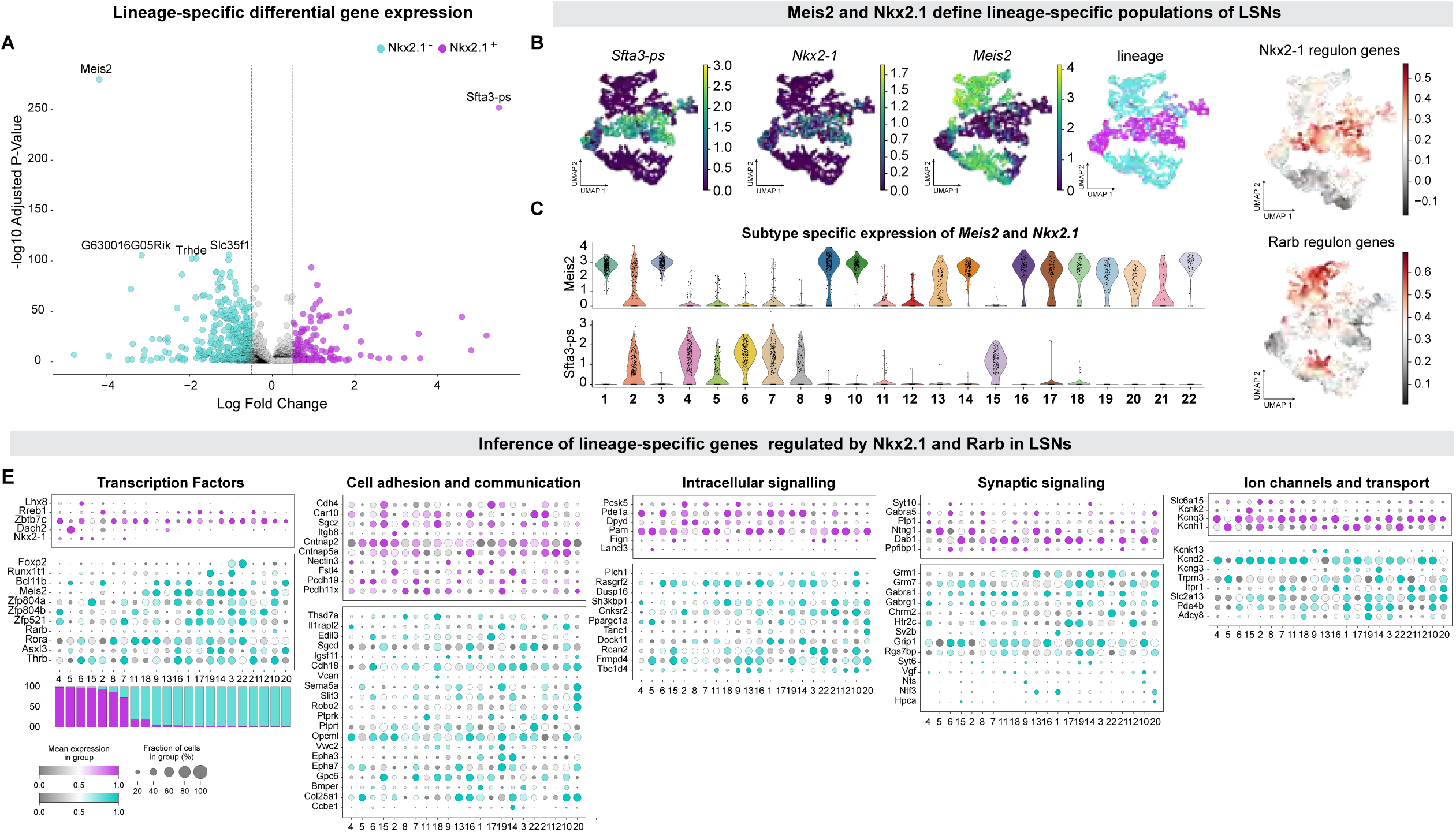
Molecular features of developmentally distinct lateral septal neurons. **A.** Volcano plot showing 684 differentially expressed genes between Nkx2.1+ (magenta) and Nkx2.1−(cyan) LSNs (157 enriched in Nkx2.1+, 527 enriched in Nkx2.1–). **B.** Feature plot showing the scaled expression (scale bar) patterns of *Sfta3-ps, Meis2,* and *Nkx2.1* among LSNs, compared with a UMAP plot of their GFP identity. **C.** Violin plots showing the scaled expression of *Sfta3-ps* and *Meis2* across the LSN subtypes. On top is a stacked bar graph showing the proportion of GFP positive (magenta) and negative (cyan) cells in each LSN subtype. **D.** Feature plots showing the module score enrichment (scale bar) of inferred target genes for NKX2.1 and RARB. **E.** Dot plot showing the scaled average gene expression (scale bar) of the top 20 differentially expressed genes between Nkx2.1+ (magenta) and Nkx2.1− (cyan) LSNs. Red asterisks mark genes that were predicted to be regulated by NKX2.1 or RARB in Nkx2.1+ and Nkx2.1− neurons respectively. On top is a stacked bar graph showing the proportion of GFP positive (magenta) and negative (cyan) cells in each LSN subtype. Subtypes are arranged by their proportion of GFP positive cells from lowest to highest.

We then explored the regulatory pathways active in LSNs using the single-cell regulatory network inference and clustering (SCENIC) computational method, allowing us to infer which transcription factor–target regulatory networks—known as regulons—were specifically active in either Nkx2.1+ or Nkx2.1− LSNs (Aibar et al., 2017). From this analysis, the NKX2.1 regulon was predicted to be specifically enriched in Nkx2.1+ LSNs, while the RARB regulon was enriched in Nkx2.1− LSNs (**Table S1**). The combined expression pattern of inferred target genes for NKX2.1 and RARB paralleled the differences in *Nkx2.1* lineage identities among the LSNs (**Figure 2D**). Several of the top differentially expressed genes distinguishing Nkx2.1+ from Nkx2.1− LSNs are predicted targets of either NKX2.1 or RARB and fall into several functional groups: transcription factors, cell adhesion and signaling, intracellular signaling, synaptic signaling, and ion channel and transport **(Figure 2E**).

### Lateral septal neuron subtypes are organized into discrete spatial domains

To spatially map LSN subtypes, we used a 500-gene panel to perform MERFISH on coronal sections of the adult septum (**Figure 3A**), including markers for medial septal neurons and non-neuronal cell types. After processing the samples, we found that a total of 263,947 cells passed our quality control filters. We then applied the same dimensionality reduction and clustering analysis as previously described (**Figure S5A**). Using established markers (**Figure S5B**), we subclustered 44,715 LSNs and integrated them with our snRNA-seq data, allowing us to identify the same 22 LSN subtypes (**Figure 3B**). When comparing the transcriptional profiles of LSN clusters between the MERFISH and snRNA-seq datasets, we observed an average cosine similarity of 0.79, indicating that they represent the same subtypes (**Figure S5D**). We profiled five sections along the anterior-posterior axis of the septum and plotted the spatial organization of each LSN subtype (**Figure 3D**), revealing that the LSN subtypes are arranged as nuclei layered in an onion-skin-like pattern, as previously described (Besnard and Leroy, 2022; Turrero García et al., 2021; Wei et al., 2012). Each subtype occupied distinct spatial domains along the three-dimensional axes of the septum. Together, these findings provide a comprehensive transcriptomic and spatial atlas of defined neuronal types in the LS.

**Figure 3:**
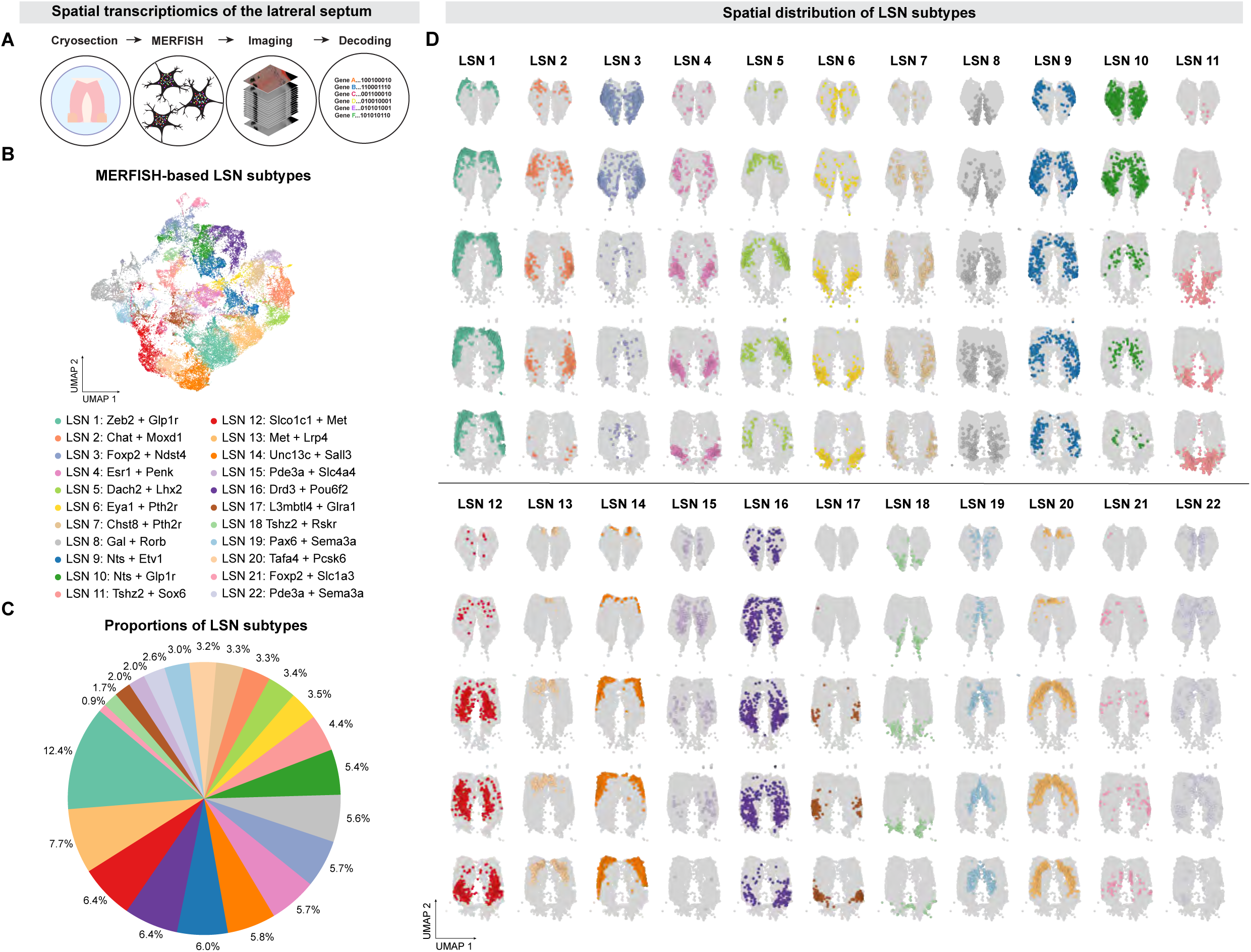
Anatomical organization of lateral septal neuron subtypes. **A.** Schematic detailing the processing of MERFISH samples. **B.** UMAP plot of 44,715 LSNs segmented from the MERFISH dataset (N=2), annotated to show LSN subtypes. **C.** Pie chart showing the percentage of total LSNs for each subtype in the MERFISH dataset. **D.** Spatial plot showing the distribution of LSN subtypes in the MERFISH dataset across 5 representative coronal sections along the anterior-posterior axis: +1.2, +0.9, +0.6, +0.3, +0.0. On top in red are the minimal marker genes necessary to identify each group.

### Spatially variable genes are a major influence on septal neuron taxonomy

Having mapped the spatial organization of LSN subtypes, we then examined the relationship between their anatomical position, molecular profile, and developmental origin. We used the statistical tool SpatialDE (Svensson et al., 2018) on our MERFISH dataset to identify 134 spatially variable genes (SVGs)—genes expressed in gradient patterns that were independent of subtype. Many SVGs shared similar patterns of gene expression and could be grouped into eight major gene sets, each characterizing a unique anatomical region of the LS: Dorsal-Lateral, Dorsal-Medial, Dorsal-Medial Inverse, Ventral, Ventral-Rostral, Ventral-Caudal, Lateral, and Medial (**Figure 4A**, **4B**).

**Figure 4:**
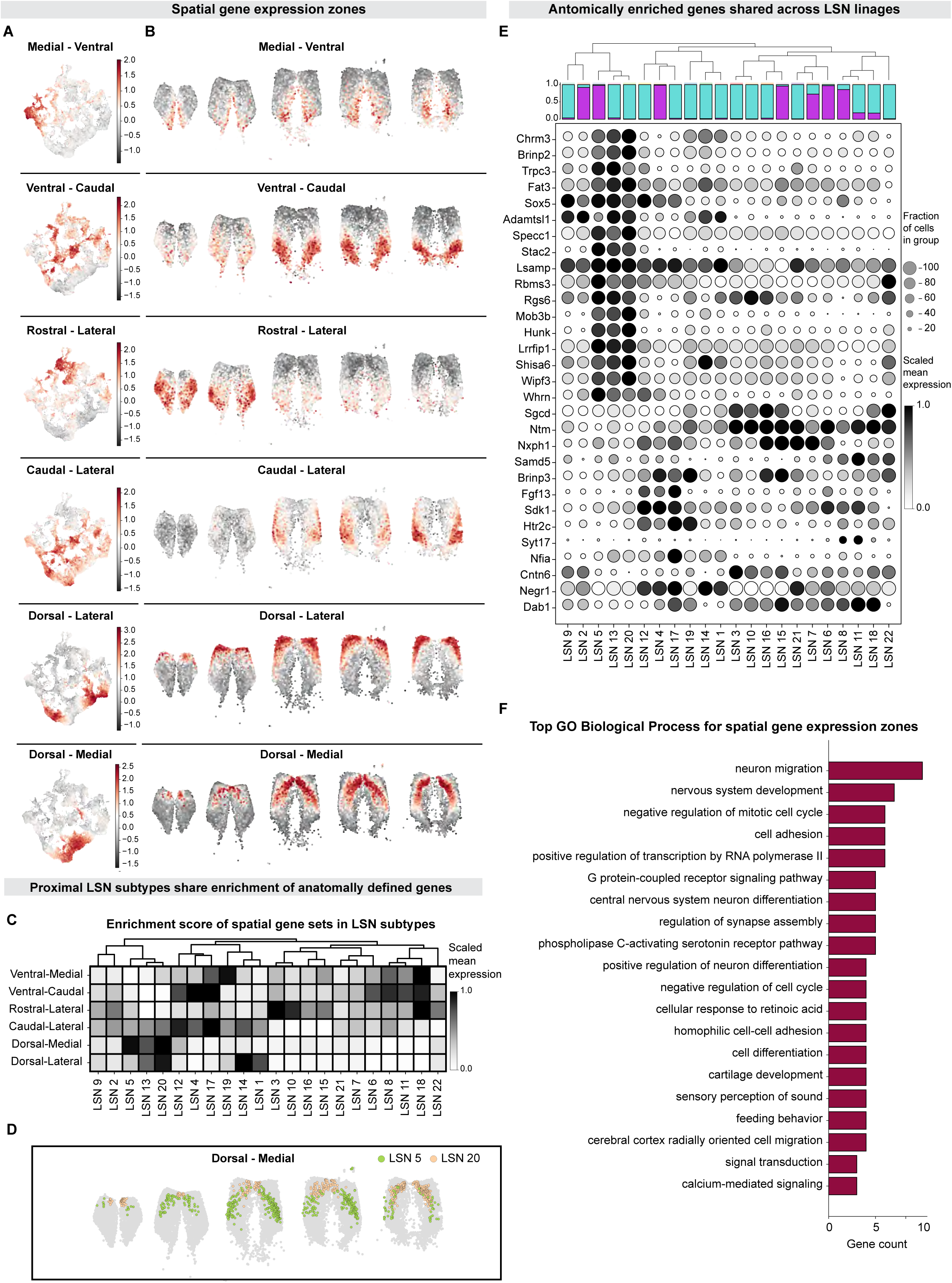
Spatially variable genes define the taxonomy of lateral septal neuron subtypes. **A.** Spatial plots showing the anatomical expression pattern for each spatially variable gene-set across 5 coronal sections of the LS and their accompanied feature plots showing the scaled module score enrichment (scale bar) for 8 gene-sets that contain spatially variable genes. **B.** Matrix plot showing the scaled score enrichment (scale bar) for each spatially variable gene-set for each LSN subtype. On top is a stacked bar graph showing the proportion of GFP positive (magenta) or negative (cyan) cells in each group, along with a dendrogram constructed from hierarchal clustering analysis using their average gene expression profiles. **C.** MERFISH spatial plots showing two pairs of subtypes that groups with similar spatial distributions and transcriptional profiles, yet come from different lineages. **D.** Dot plot showing the scale gene expression of genes within the spatially variable modules. **E.** Bar plot constructed from gene ontology analysis showing the number of spatially variable genes associated with a given molecular function. The scale bar shows the adjusted p-value for each molecular function

Notably, the expression profile of these SVG patterns across the LSN subtypes largely reflected the hierarchical relationships of their average gene expression (**Figure 4C**). For example, LSN-5, LSN-13, and LSN-20 have enriched expression of Dorsal-Medial genes, whereas LSN-4, LSN-12, and LSN-17 primarily express Ventral-Caudal genes. These patterns suggest that for many subtypes, anatomical location, rather than lineage origin, is a primary driver of transcriptional similarity among LSNs—as exemplified by LSN-5 and LSN-20, which reside in overlapping domains yet derive from Nkx2.1+ and Nkx2.1− lineages, respectively (**Figure 4D, E**). We performed GO analysis to investigate the functions of SVGs and found that many are involved in neuronal migration, synaptic development, and cell adhesion (**Figure 4F**). In conclusion, these findings demonstrate that the expression of SVGs can shape the molecular identity of LSNs independently of lineage origin.

### Molecularly defined lateral septal neuron subtypes exhibit distinct electrophysiological, morphological, and anatomical features

Early classifications of LSNs were based on their physiological and morphological properties (Alonso and Frotscher, 1989; Gallagher et al., 1995b; Raisman, 1969), yet how these characteristics correlate with their molecular identity remains unclear. To investigate this, we leveraged our sequencing data to identify marker genes with available mouse Cre lines that label non-overlapping subgroups of LSNs (**Figure 5A**). We first validated this approach by injecting Foxp2-Cre and Chat-Cre animals with a Cre-dependent AAV expressing nuclear-localized yellow fluorescent protein and performing snRNA-seq on sorted fluorescent-positive septal nuclei (**Figure S6 and B**). In the Foxp2-Cre line, GFP-positive nuclei were predominantly within clusters LSN-3 and LSN-22, both of which selectively express *Foxp2*. Similarly, in the Chat-Cre line, GFP-positive nuclei were predominantly enriched in LSN-2, which selectively expresses *Chat*. Having validated this marker-selection approach, we used viral methods to label neurons from four Cre lines (Esr1, Foxp2, Lhx2, and Ndnf) and performed whole-cell patch clamp recordings followed by neurobiotin filling to characterize their intrinsic physiological and morphological properties. Each subgroup exhibited distinct firing types and action potential waveforms. Using 14 intrinsic parameters collected from the recordings, we compared the electrophysiological profile of each subgroup and found that *Esr1* and *Foxp2* neurons shared similar intrinsic properties, while *Lhx2* neurons were the most distinct (**Figure 5D**).

**Figure 5:**
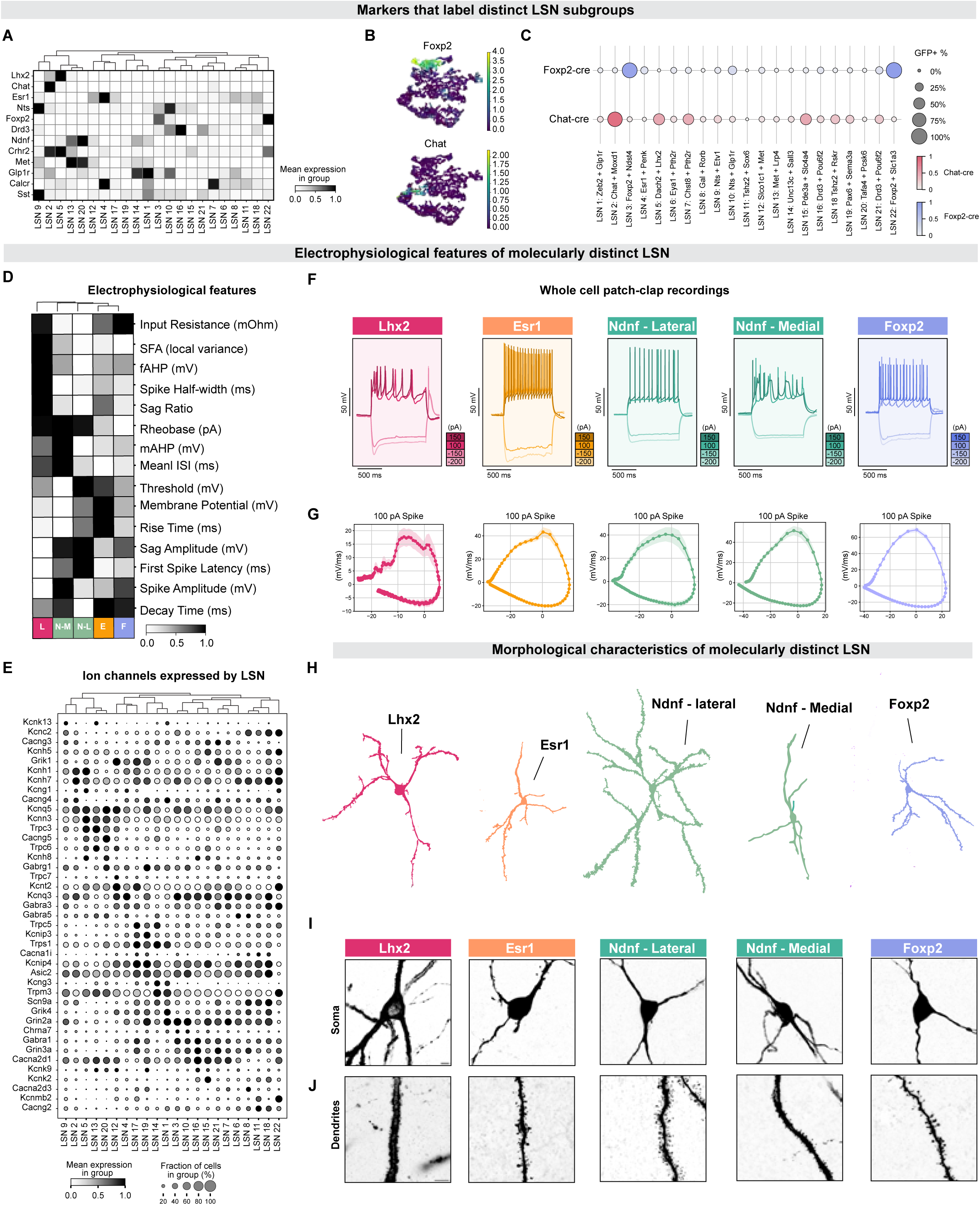
Electrophysiological and morphological properties of lateral septal neuron subgroups. **A.** Matrix plot showing the scaled average expression across the LSN subtypes of marker genes with available Cre lines. **B.** Dot plot of *Foxp2* and *Chat* expression of sorted nuclei from Cre lines that mark non-overlapping subgroups of LSNs. **C.** Bar plot showing the proportion of cells labeled within Chat-Cre (top, Blue) line and Foxp2-Cre (bottom, Red) line. **D.** Morphological traces from neurobiotin filled LSNs for each genetically defined group. **E.** High magnification images of neurobiotin filled (Esr1, Ndnf-Lateral, Ndnf-Medial, and Foxp2) or tdTomato fluorescent (Lhx2) cells for each genetically defined group. The top row shows their somas, and bottom bar shows their dendrites (scale bars 10 μm). **F.** Representative traces showing the firing pattern of cells recorded using voltage clamp from each genetically defined LSN subgroup. Current was injected in 25pA steps ranging from −200 to 150pA, with higher currents shaded in darker colors. **G.** Phase plot showing the membrane voltage rate change for a representative action potential for a 100pA current injection for each LSN subgroup. **H.** Heatmap showing the scaled values for 14 intrinsic properties. On top is a dendrogram showing the relationships among subgroups based on the intrinsic properties.

The firing patterns of each population varied from each other when injected with a series of current steps (**Figure 5D–F**). *Ndnf* neurons showed two distinct firing patterns and morphologies dependent on their topographical location along the medial-lateral axis of the LSd. This separation corresponded to the anatomical distribution of LSN-4 (lateral) and LSN-6 (medial). At high current injections, *Esr1*, *Foxp2*, and lateral *Ndnf* neurons fired and sustained a regular sequence of action potentials. In contrast, *Lhx2* and medial *Ndnf* neurons spiked irregularly, each showing a unique firing pattern. *Lhx2* neurons spiked transiently, producing only a few action potentials that had successive reductions in peak amplitude and widening of the spike width. When subjected to high current injections, medial *Ndnf* neurons displayed an erratic firing pattern characterized by a combination of regular spikes and plateau potentials. All groups except for *Ndnf* neurons demonstrated a prominent inward-hyperpolarizing current at negative current injections, a characteristic of inward-rectifying potassium channels (Hibino et al., 2010; Mao et al., 2003). The electrical properties of neurons are determined by the complement of voltage-gated ion channels that they express (Bean, 2007; Burke and Bender, 2019); using our snRNA-seq dataset, we identified sets of voltage-gated ion channels that were differentially expressed among LSNs. In line with their intrinsic physiological relationships, *Esr1* and *Foxp2* shared a similar enrichment of channels. Notably, *Lhx2* neurons expressed the largest number of uniquely expressed channels, primarily from the voltage-gated potassium channel family (**Figure 5E**). Lastly, we visualized recorded neurons by staining for neurobiotin with fluorescently labeled streptavidin. Each subgroup displayed distinct morphological features, including differences in soma size, dendritic arborization, and spine density (**Figure 5H-I**).

### Lateral septal neuron subtypes target distinct output regions and receive subtype-specific inputs

We then sought to systematically map the connectivity of molecularly distinct LSN populations. The efferent projections of each subgroup were labeled by local injection of an AAV expressing a Cre-dependent eGFP and synaptophysin-fused mRuby, into the LS of each of the four transgenic lines (for more specific labeling, we opted to use the Chat-Cre line—which primarily labels LSN-5—instead of the Lhx2-CreER line) (**Figure 6A-C**). The fibers of each subgroup displayed unique projection patterns, targeting distinct hypothalamic, subpallial, and midbrain regions (**Figure 6D, S7A**).

**Figure 6:**
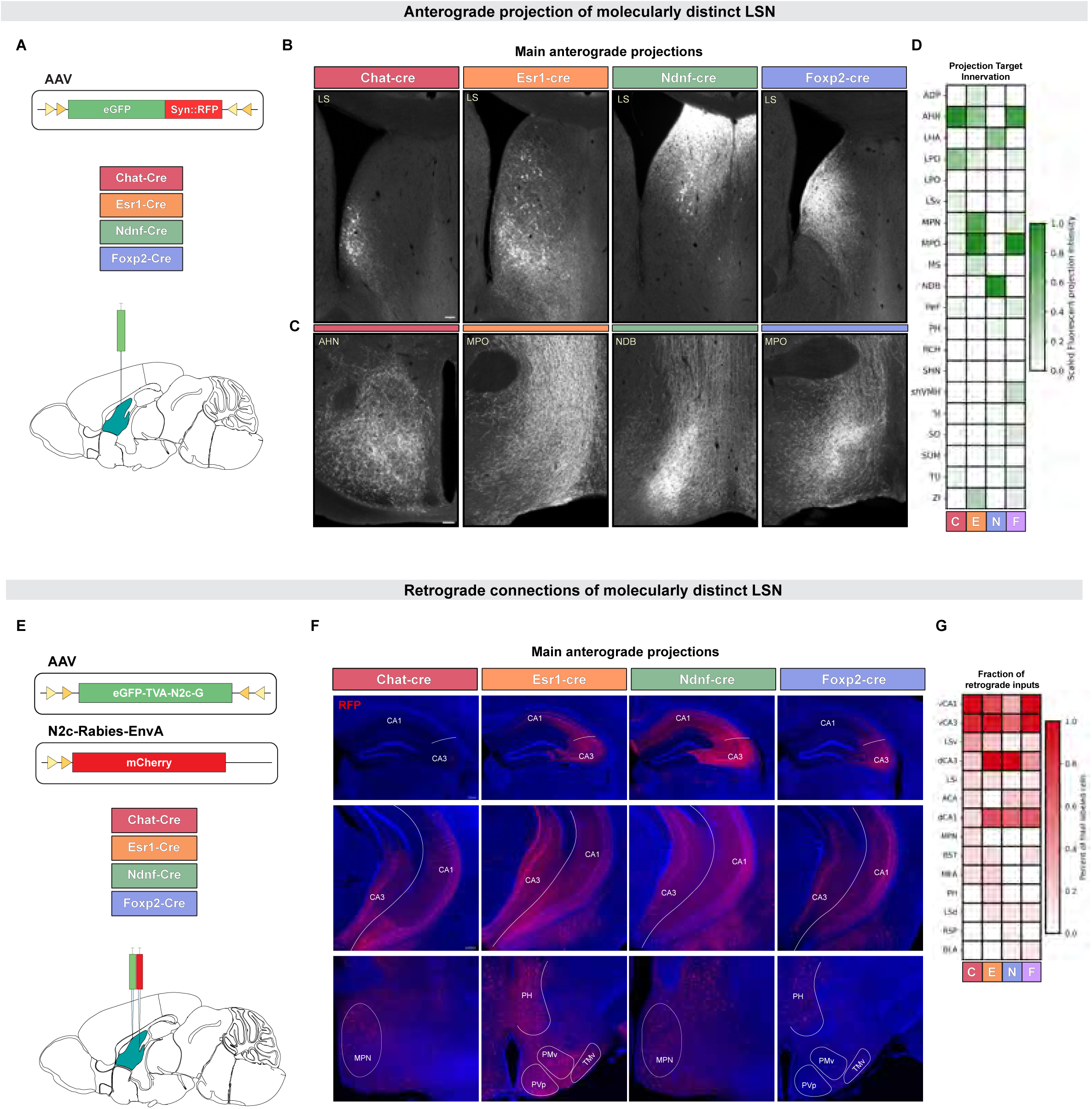
Anatomical connections of lateral septal neurons subgroups. **A.** Schematic showing a local injection of an AVV expressing a Cre dependent eGFP and synaptophysin fused RFP into the LS of four transgenic. **B.** Representative images for each transgenic line showing coronal sections of the LS immunofluorescently stained for GFP after a local AAV. Scale bar: 100 μm. **C.** Representative images showing GFP-expressing projections of LSN subgroups labeled in each Cre line. Scale bar: 100 μm. **D.** Matrix plot illustrating the regional proportion of total synaptophysin fluorescence for each Cre line: Chat-Cre (N = 4), Esr1-Cre (N=3), Ndnf-Cre (N=4), Foxp2-Cre (N=2). **E.** Schematic showing a helper AAV expressing a Cre-dependent eGFP, TVA, and N2c-G protein, along with the Rabies-N2c virus being injected into the LS of each transgenic line. **F.** Representative images for each transgenic line showing mono-synaptically labeled cells labeled with a tdTomato expression rabies-N2c. Matrix plot illustrating the regional proportion of total Rabies labeled cells foreach Cre line: Chat-Cre (N = 3), Esr1-Cre (N=2), Ndnf-Cre (N=4), Foxp2-Cre (N=2).

*Esr1* and *Foxp2* neurons exhibited distributed projection patterns targeting various hypothalamic areas, with no single area receiving more than 25% of their total projections. The terminals of both groups highly innervated the medial preoptic area (MPO), a key regulator of physiological states and social behaviors (Dulac et al., 2014). In contrast, *Chat* and *Ndnf* neurons showed more restricted projection patterns, with one or two regions accounting for the majority of the total projections. The terminals of *Chat* neurons densely innervated the anterior hypothalamic area (AHN), while *Ndnf* neurons selectively targeted the nucleus of the diagonal band (NDB) and the lateral hypothalamic area (LHA) (**Figure 6D**). Chat neurons (LSN-2) are a subset of *Crhr2*-expressing LSNs, whose projections to the AHN have been shown to regulate threat-response behaviors (Anthony et al., 2014; Hashimoto et al., 2022). The NDB and LHA are key modulators of cognitive processes, including attention, learning, goal-seeking and memory consolidation (Yamashita and Yamanaka, 2016; Petrovich, 2018; Liu et al., 2018). Collectively, these findings reveal significant differences in the postsynaptic targets of molecularly distinct LSNs, likely reflective of the behaviors they modulate.

We then used mono-synaptic rabies tracing in combination with the same transgenic Cre lines to identify presynaptic inputs onto each genetically defined group (**Figure 6E, S7B**). While each group received inputs from numerous brain regions, the hippocampal and hypothalamic subregions contributed the largest proportion (**Figure 6F**), consistent with previous reports (Albert et al., 1978; Gergues et al., 2020; Raisman, 1969; Risold and Swanson, 1997). The majority of hippocampal inputs onto *Chat* neurons came from the ventral CA1 and CA3 subregions. Similarly, *Foxp2* neurons predominantly received innervation from ventral hippocampus regions, but also received inputs from its dorsal regions. Interestingly, *Esr1* neurons received an even distribution of inputs from the dorsal and ventral hippocampal regions with a higher contribution from the CA3 region than CA1 (28.4% vs. 16.1%). *Ndnf* neurons had the largest proportion of input from the dorsal hippocampus, mostly from CA3 areas. Together, these data reveal that molecularly defined LSN subtypes occupy distinct positions within the limbic circuit, with subtype-specific input–output relationships that likely underlie their functional specialization.

## DISCUSSION

Neurons in the lateral septum (LS) regulate complex affective, social, and motivational processes, and are organized topographically along its anatomical axis (Besnard et al., 2019; Menon et al., 2022; Sheehan et al., 2004; Wirtshafter and Wilson, 2021). A complete understanding of their specialized functional properties requires characterization across multiple modalities (Gouwens et al., 2020; Peng et al., 2021; Wang et al., 2023; Zeng, 2022; Zhang et al., 2023). Some of these features are intrinsically encoded by lineage-related developmental programs, while others are shaped by extrinsic factors from the local environment. Here, we resolve the multimodal features and organizational principles of lateral septal neurons (LSNs) by detailing their transcriptome, developmental origin, synaptic partners, and physiological properties, identifying 22 molecularly distinct LSN types shaped by both developmental lineage and anatomical location.

Nkx2.1+ LSNs comprise at least four distinct subtypes that share a common transcriptional signature, most prominently the expression of lncRNA Sfta3-ps (also known as NANCI). Sfta3-ps lies adjacent to Nkx2.1 in the genome and is thought to positively regulate its expression (Herriges et al., 2017, 2014; Katayama et al., 2005), suggesting that it helps maintain the low Nkx2.1 expression we observe in mature LSNs — in contrast to cortical interneurons, which downregulate Nkx2.1 post-mitotically (Elias et al., 2008). This postmitotic maintenance of Nkx2.1 expression mirrors that of basal ganglia neurons, where Nkx2.1 contributes to proper migration and wiring (Magno et al., 2017; Nóbrega-Pereira et al., 2008). Additionally, the expression of Nkx2.1 in LSNs may repress molecular programs enriched in neurons derived from the lateral ganglionic eminence (LGE) as observed when Nkx2.1 is ablated in MGE-derived cortical interneurons (Sandberg et al., 2016).

In line with this idea, Nkx2.1− neurons selectively express Meis2, a TALE-homeodomain transcription factor highly expressed in the LGE that is required for the differentiation of both D1 and D2 medium spiny neurons (Su et al., 2022). Consistent with an LGE-like transcriptional state, many Nkx2.1− LSNs co-express Bcl11b and Runx1t1, regulators of MSN differentiation (Arlotta et al., 2008). Our SCENIC analysis predicted RARB as regulator of these factors, a factor known to control the development of striatonigral projection neurons (Rataj-Baniowska et al., 2015), suggesting that retinoic acid signaling further promotes the LGE-like identity in this population. Together, these transcriptional programs indicate that Nkx2.1− LSNs retain a molecular signature reminiscent of LGE-derived striatal neurons, with Meis2 acting in concert with Bcl11b, Runx1t1, and retinoic acid signaling to specify and consolidate this identity.

Using MERFISH, we generated a comprehensive spatial atlas, detailing the discrete arrangement of subtypes within a layered pattern from the midline, consistent with the onion-skin organization of the septum (Wei et al., 2012). Several progenitor-domain transcription factors — Pax6, Lhx2, Zeb2, and Nkx2.1 — remain expressed in anatomically separated and molecularly distinct subtypes of LSNs, suggesting that these transcription factors may play a role in establishing the cytoarchitecture of the lateral septum (Flames et al., 2007). Clarifying how these developmental domains establish adult topography will require detailed clonal mapping (Bandler et al., 2022; Delgado et al., 2022; Harwell et al., 2015; Mayer et al., 2015).

In addition to their lineage features, the molecular identity of LSNs was also defined by sets of spatially genes expressed within defined anatomical domains, driving Nkx2.1+ and Nkx2.1− cells adopt similar transcriptional states. LSN-5 and 20 subtypes contain cells from predominantly Nkx2.1+ and Nkx2.1−lineages respectively, and express and are co-enriched for an array of features, many of which play a role in synaptic function, such as Shisa6, Rbsm3, and Hunk. These SVG may be reflective of the organization of hippocampal inputs into the LS (Besnard and Leroy, 2022; Risold and Swanson, 1997).

We recorded from four non-overlapping LSN subgroups and found that their firing patterns fell broadly into two classes: regular-spiking (Foxp2/LSN-2, 5, Esr1/LSN-5, and lateral Ndnf/LSN-1, 14) and irregular-spiking (Lhx2/LSN-8 and −11; medial Ndnf/LSN-13 and 20). The prevalence of regular, non-adapting firing is consistent with a role in relaying continuous signals from the hippocampus, particularly for dorsally located cells that encode locomotion and spatial context (Bender et al., 2015; Besnard et al., 2019; Wirtshafter and Wilson, 2020, 2021). The transient firing of Lhx2 neurons is compatible with expression of the SK-type channel Kcnn3 (Sivaramakrishnan and Oliver, 2001), while the plateau potentials of medial Ndnf neurons likely involve voltage-gated calcium or persistent sodium currents (Teka et al., 2011). Irregular-spiking subgroups project to regions implicated in motivation, attention, and threat response (lateral hypothalamus, nucleus of the diagonal band, and anterior hypothalamic area), and share a stubby-spine morphology associated with distinct synaptic properties compared to mushroom spines (Hayashi and Majewska, 2005; Pchitskaya and Bezprozvanny, 2020).

Rabies tracing and anterograde labeling showed that the hippocampus is the dominant input for each subgroup we examined, and the hypothalamus their dominant target (Albert et al., 1978; Gergues et al., 2020; Raisman, 1969; Risold and Swanson, 1997), however input–output distributions vary by subtype. Ndnf neurons were the only subgroup to receive predominantly dorsal hippocampal input and to preferentially target the LHA and NDB — a pattern previously described for dorsolateral Sst+ LSNs (Besnard et al., 2019), likely corresponding to LSN-4a. Because Ndnf labels several subtypes we cannot fully disambiguate their connectivity, raising the possibility that these two subtypes innervate overlapping gross targets but distinct postsynaptic cell types. A similar scenario may apply to LSN-1 and LSN-9, which share Nts expression and occupy adjacent rostral domains and are likely to contribute to feeding and energy-balance circuits (Azevedo et al., 2020; Chen et al., 2022; Li et al., 2023). Together, these observations highlight opportunities for more selective intersectional tools to resolve the circuitry of individual subtypes.

The identity and function of LS neurons are shaped by intrinsic lineage programs and further refined by extrinsic spatial and experience-dependent factors. Our data support the view that heterogeneous LSN subtypes operate collectively as functional units to regulate affective, motivational, and physiological processes (Besnard and Leroy, 2022). The multimodal atlas presented here provides a foundation for dissecting LS circuit function in health and disease.

### Limitations of the study

Several limitations should be considered when interpreting our results. First, the molecular relationships among LSN subtypes were summarized with hierarchical clustering, which imposes a binary tree structure. UMAP projections suggest more continuous relationships among Nkx2.1−lineage subtypes, and alternative representations — such as constellation plots (Yao et al., 2023) — may capture inter-type relationships more faithfully; claims that depend on tree position should be interpreted within this caveat. Our functional characterization relied on transgenic lines selected to label non-overlapping subgroups, but the correspondence between driver lines and transcriptomic subtypes was inferred from marker-gene expression rather than confirmed by snRNA-seq profiling of labeled cells. Some driver lines (notably Lhx2-Cre, which captures both LSN-8 and LSN-11; and Ndnf-Cre, which captures both LSN-4 and LSN-6) therefore aggregate subtypes we cannot further resolve in the physiological and anatomical data. Morphological and electrophysiological differences across subgroups are illustrated with representative cells; population-level quantification of spine density, soma size, and dendritic thickness remains a future direction. The snRNA-seq comparisons relied on FACS sorting of GFP+ versus GFP– nuclei. While we observed no evidence of transcriptional saturation or systematic artifacts attributable to GFP expression, non-specific effects on the mature transcriptome cannot be fully excluded. The snRNA-seq and MERFISH datasets were generated from mice aged ∼5 weeks, whereas electrophysiological and circuit tracing experiments used mice aged 2–6 months to accommodate stereotaxic surgery and viral expression time courses. Although LS transcriptomic and spatial organization is expected to be stable across this interval, we cannot rule out age-related variability contributing to the functional data, particularly given the modest sample sizes. Finally, the MERFISH-to-snRNA-seq mapping relies on a 500-gene panel and identifies some cross-correlations (e.g., between LSN-5 and LSN-7) that are higher than expected given their separation in other analyses. These likely reflect limits of the chosen gene panel rather than true transcriptomic equivalence.

## KEY RESOURCES TABLE

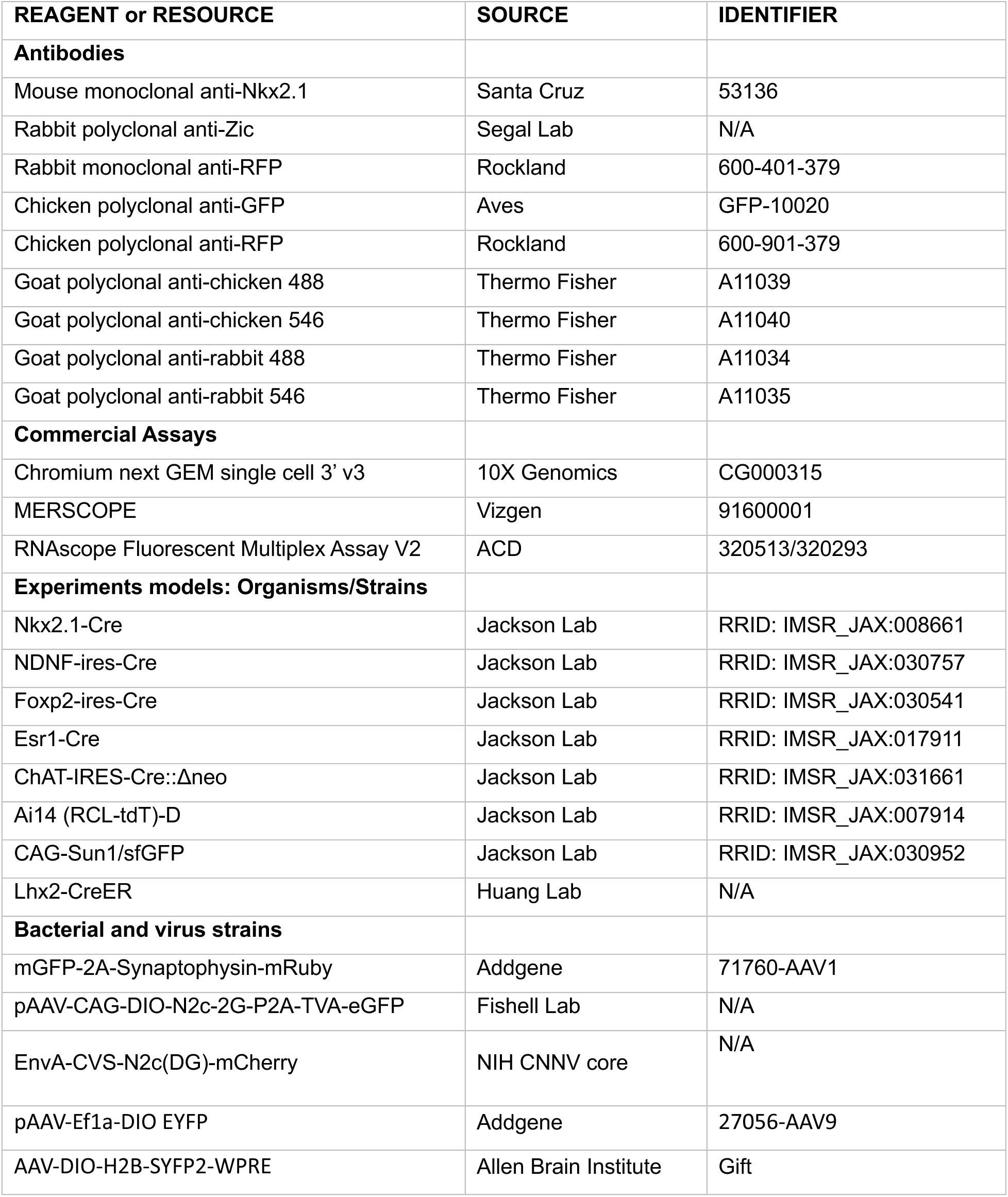

## LEAD CONTACT AND MATERAL AVAILABILITY

No new reagents were produced from this study. Further information and requests for resources and reagents should be directed to and will be fulfilled by the lead contact, Corey Harwell

## EXPERIMENTAL MODEL AND STUDY PARTICIPANT DETAILS

### Animals

All animal procedures were reviewed and approved by the Institutional Animal Care and Use Committee (IACUC) at both Harvard University and the University of California, San Francisco. The welfare and care of the animals, as well as the conduct of the experiments, strictly adhered to the NIH guidelines. The present study was conducted utilizing both male and female adult mice of C57Bl6/J genetic background, with ages ranging between 30 and 90 days. Animal housing conditions adhered to a 12-hour light/dark cycle, with food and water supplied ad libitum.

## METHOD DETAILS

### Immunofluorescence

Adult animals were anesthetized with 2.5% Avertin, and were transcardially perfused with PBS, followed by 4% PFA in PBS. Subsequently, their brains were dissected out and subjected to post-fixation in 4% PFA at 4°C overnight. Brain tissues were then sectioned into slices of 75–100 µm thickness using a vibratome (Leica Microsystems VT12000S) and preserved in a freezing buffer (70 g sucrose, 75 ml ethylene glycol, filled to 250 ml with 0.1 sodium phosphate buffer) at −20°C. Before further processing, the sections underwent a washing cycle, consisting of three 5-minute washes in PBS. The sections were then permeabilized in a solution of 10% serum and 0.3% Triton in PBS for one hour. The primary antibodies were diluted in the same solution at concentrations ranging between 1:500-1:1000, and the sections were incubated in this for 24 hours at 4°C. Following another round of washing, as performed previously, the sections were incubated in secondary antibodies, which were diluted in 10% serum in PBS at a concentration of 1:1000. This incubation step lasted for 2 hours at room temperature. After one final wash, the sections were labeled with DAPI (4’,6-diamidino-phenylindole, Invitrogen) and subsequently mounted onto a glass slide and cover-slipped using an antifade mounting medium.

### Single-nucleus RNA-sequencing and library preparation

A total of six mice (3 male and 3 female) across three replicates were used for this study. Nkx2.1−Cre mice crossed with the conditional Sun1GFP allele were 35 days old at the time of sacrifice. The brain tissue was carefully extracted, and the septum was selectively dissected and immersed in a hibernation buffer consisting of 0.25M sucrose, 25mM KCl, 5mM MgCl2, and 20mM Tricine KOH. Following this, the dissected tissue was gently transferred into a pre-chilled 2 ml Dounce homogenizer containing 500 µl of hibernation buffer supplemented with 5% IGEPAL, half a Roche protease inhibitor tablet, 0.2 U/µl Promega RNasin, 0.1% spermidine, 0.1% DTT, and 0.1% spermine. The tissue was delicately homogenized using loose and tight pestles, approximately 15 strokes each. After a resting period of 10 minutes, the homogenate was transferred to a low-bind Eppendorf tube and centrifuged for 5 minutes at 500 g at 4 °C. The supernatant was discarded, and the pellet was carefully resuspended in 500 µl of 1% BSA in PBS supplemented with 0.2 U/µl Promega RNasin, 0.1% spermidine, 0.1% DTT, and 0.1% spermine. The suspension was centrifuged once again under similar conditions, and the pellet was resuspended as previously outlined. The final suspension was strained through a 40 μm Flowmi filter and stained with 0.8 µl of Ruby Dye prior to Fluorescence-activated Cell Sorting (FACS). Nuclei double positive for GFP and Ruby Dye were separated from nuclei only positive Ruby dye using a SONY SH800S FAC sorter. After collecting a minimum of 50,000 events, the nuclei were loaded onto a 10X Chromium Single Cell 3’ chip (v3 chemistry) at the specified concentration. This was followed by adherence to the standard 10X Chromium Single Cell 3’ protocols (v3 chemistry). Finally, the resultant libraries were sequenced using a NovaSeq 6000 S4 flow cell, achieving a read depth of approximately 50,000 reads per nucleus.

### Single nuclei-sequencing analysis

The sequenced reads were aligned to the mouse genome (mm10), inclusive of intronic reads, using the Cellranger 7.1.0 software package from 10X Genomics. The mean number of genes per cell was 3,002. For the subsequent analysis, we utilized the Python module Scanpy (Wolf et al., 2018). The raw count matrix of each sample was loaded into a AnnData file, and all samples were concatenated into one AnnData file. Genes expressed in less than 3 cells were filtered out, along with cells that expressed fewer than 200 genes.

We then normalized the raw counts to 10,000 reads per cell before identifying the highly variable genes. Following this process, we performed principal component (PC) analysis and computed the PC variance ratio to determine how much each PC contributed to the total variance. From this, we determined that the first 15 PCs were sufficient to explain the total variance and were used to construct a neighborhood graph of our cells. Subsequently, this graph was embedded in two-dimensions for visualization using unifold manifold approximation and projection (UMAP) dimensionality reduction. Batch effect between replicates was corrected for using BBKNN (Polański et al., 2020). We then performed Leiden clustering at multiple resolutions to understand how the clusters evolved as a function of resolution. Using a resolution of 0.9 we performed pairwise differential gene analysis between the clusters and all other cells using the Wilcoxon rank-sum test. Our general cell types were annotated by cross-referencing established marker genes or evaluating the regionalization of genes enriched in our clusters that were accessible via the Allen Brain *in situ* hydrization resource (Lein et al., 2007). Once general cell types were established, we subclustered the LS neurons and performed the same dimensionality reduction, embedding, and marker gene analysis mentioned previously.

### SCENIC Regulon analysis

Regulon analysis was performed using the pySCENIC Python implimintation of the SCENIC pipeline (Aibar et al., 2017). The raw count matrix for LS neurons was used to ‘infer regulons (transcription factors and their targets) based on correlations of gene expression across cells using the arboreto package, which utilizes GRNBoost2 (Moerman et al., 2019). Target genes that had no enrichment for the transcription factor binding motif within a genome search space between 500 bp and 10kb around the transcriptions start site were then pruned. Subsequently, the ‘aucell’ package was used to compute an area under curve value for each regulon across all cells. Finally, regulon specificity scores were computed for both GFP-positive and -negative LS neurons to determine which regulons were enriched in either population.

### Gene ontology enrichment

We employed the ‘ClusterProfiler’ R package to conduct gene ontology (GO) analysis on GFP-positive and -negative LS neurons (Wu et al., 2021; Yu, 2020). Initially, differentially expressed genes between these two types of neurons were identified. Genes co-expressed in clusters with a majority of GFP-positive or -negative cells were then selected. The resultant gene list was subjected to a hypergeometric test to pinpoint overrepresented GO terms, categorized under three domains: biological process, cellular component, and molecular function. The same process was applied to the list of spatially variable genes. Terms with a p-value below 0.05 were deemed significantly enriched.

### Selection of MERFISH Gene Panel

A multi-pronged approach was used to curate a 500 gene MERFISH panel, aiming to identify the same transcriptionally distinct cell types found in the single-nucleus RNA-seq dataset. We began by manually incorporating genes recognized as well-established general cell type markers. Subsequently, used the single-nucleus sequencing dataset as reference to identify genes showing enrichment in each of our cell types of interest. This was accomplished by performing pairwise differential gene expression analysis between cell groups. Finally, we employed the random forest classifier scRFE on the single-nucleus dataset to identify the sets of genes that best describe our cell groups (Park et al., 2020). This list was uploaded and approved for probe encoding by the Vizgen Gene Panel Design Portal.

### MERFISH sample preparation and hybridization

We utilized 35-day-old C57Bl6/J mice, both male and female, in our study. Following anesthesia with 2.5% Avertin, the animals were scarified and their brain tissues were carefully extracted. The harvested tissues were immediately preserved in OCT (Optimal Cutting Temperature compound) and uniformly frozen using dry ice. Subsequently, the tissues were stored at −80°C for future use. 24 hours prior to sectioning, the samples were warmed to −20°C, after which serial cryosectioning was conducted at a thickness of 10 micrometers. Several coronal sections, representing different areas of the LS along the rostral-caudal axis, were mounted onto a glass slide from the Vizgen MERSCOPE Slide Box. The mounted sections were fixed with 4% PFA in 1X PBS, rinsed, permeabilized, and preserved in sterile 70% ethanol. The procedures of probe hybridization, gel-embedding, and tissue clearing were performed according to the MERSCOPE sample preparation user guide. Briefly, we washed the sections with a formamide wash buffer (PN 20300003) and incubated them for 30 minutes at 37°C, before the probe hybridization mix was added. The sections were then incubated at 37°C for 48 hours while submerged in the MERFISH library mix. After probe incubation, we washed the sections with formamide buffer two times for 30 minutes each at 47°C, and once with sample prep wash buffer (PN 20300001). We then applied a gel embedding solution containing 10% ammonium persulfate, tetramethylethylenediamine, and a gel embedding premix (PN 20300004). A cover slip provided by the MERSCOPE Slide Box was used to slowly spread the gel embedding solution over the sections, and the gel was solidified at room temperature for 1.5 hours. Once solidified the coverslip was slowly removed, and a clearing solution containing Warm Clearing Premix (PN 20300003) and proteinase K was incubated with the sections at 37°C for 24 hours.

### MERFISH image analysis and cell segmentation

After gel clearing, the samples were rinsed with the Wash Buffer provided in the Vizgen Imaging Kit and then placed into the MERSCOPE instrument flow cell. After the assembly of fluidics and requisite reagents, a low-resolution image of DAPI signal was captured using a 10X objective lens in a tilescan format. Areas exhibiting well-preserved lateral septal samples were delineated as regions of interest. Following the marking of all such regions, multiple rounds of 3-color imaging were conducted using a 60X objective lens. During each imaging round, seven focal planes along the z-axis were captured for each channel. The raw image files obtained were processed via the MERlin image analysis pipeline (Moffitt et al., 2018). The Cellpose2 software was utilized to segment cells from the nuclear DAPI signal (Pachitariu and Stringer, 2022). The decoded RNA molecules were then partitioned into individual cells to generate single-cell count matrices.

### MERFISH Clustering Analysis

The coordinates and count matrix from the MERFISH data were imported as an AnnData object. Cells with a volume larger than 200 um^^03^, with fewer than 100 transcripts, and with a DAPI score less than 500 were excluded from the analysis. The remaining cells were then subjected to standard single-cell analysis following previously described protocols.

### Spatially variable gene patterns

Spatially variable genes were ascertained from our MERFISH data utilizing SpatialDE (Svensson et al., 2018). This analysis allowed us to model gene expression levels in relation to the spatial coordinates of cells and assign rankings to genes based on the degree of spatial variability in their expression. Following this, automatic expression histology was employed to recognize genes exhibiting similar patterns of expression across the tissue. This methodology was applied to five representative sections. By manually matching patterns across these sections and annotating them based on their expression throughout the lateral septum, we were able to generate a detailed map of spatially variable gene expression.

### Stereotaxic Viral Injections

All surgical procedures were executed under aseptic conditions. Mice aged 2-6 months were anesthetized with isoflurane, followed by an injection of either 1.5 mg/kg SR Buprenorphine or 3.25 mg/kg EthicaXR, in addition to a 5 mg/kg dose of Meloxicam. They were then placed on a heating pad with their head fixed on a stereotaxic apparatus (Kopf). Their eyes were protected from drying using an ophthalmic ointment, and their hair was removed from the scalp to allow for an incision that exposed the cranium. After leveling the skull, a hole was drilled into it at coordinates that targeted the lateral septum: ChATCre: AP +0.49 mm, ML 0.47 mm, DV −3.3 mm; Esr1Cre: AP +0.25 mm, ML 0.5 mm, DV −3.3 mm; NdnfCre: AP +0.13 mm, ML 0.3 mm, DV −2.5 mm; Foxp2Cre: AP +1.09 mm, ML 0.45 mm, DV −3.5 mm. Glass capillaries were backfilled with mineral oil and front-filled virus, before being loading into a Hamilton syringe. Subsequently the syringe was connected to a nanoliter injector (WPI) which delivered our targeted volume of injection at a rate of 25 nl/s. After 10 minutes, the capillary was slowly removed from the brain and the incision was sutured. The mice were allowed to recover partially from anesthesia before being returned to their cage.

For the anterograde tracing experiments, 50-100 nl of pAAV hSyn FLEx mGFP-2A-Synaptophysin-mRuby was injected into the lateral septum of Ndnf-Cre, Esr1-Cre, Chat-Cre, and Foxp2-Cre mice. The monosynaptic rabies tracing experiments required two surgical procedures. First mice were injected with 50-100 nl of a helper virus pAAV-CAG-DIO-N2c-2G-P2A-TVA-eGFP. Three weeks later a second surgery was performed to inject 50 nl of the Rabies EnvA-CVS-N2c(DG)-mCherry virus. The mice were euthanized five days after the second injection, and their brain tissue was collected.

### Electrophysiology recordings and cell filling

Adult mice were initially anesthetized with 2.5% Avertin, followed by transcardial perfusion with 5 ml of ice-cold, oxygenated slice solution saturated with 95% O2 and 5% CO2. The slice solution consisted of the following components (in mM): 110 choline chloride, 2.5 KCl, 0.5 CaCl2, 7 MgCl2, 1.3 NaH2PO4, 25 NaHCO3, 10 glucose, 1.3 Na-ascorbate, and 0.6 Na-pyruvate. The osmolarity of the slice solution was adjusted to 305–315 mOsm/L using sucrose. After perfusion, the brains of the mice were extracted and placed in ice-cold, oxygenated slice solution. Using a vibratome (VT1200s, Leica), the brain tissue was carefully sectioned into 200 µm thick slices. These slices were then incubated for at least 1 hour at 33°C in oxygenated artificial cerebrospinal fluid (ACSF) containing (in mM): 125 NaCl, 2.5 KCl, 2 CaCl2, 1.3 MgCl2, 1.3 NaH2PO4, 1.3 Na-ascorbate, 0.6 Na-pyruvate, 10 glucose, and 25 NaHCO3 (305-315 mOsm/L). Following incubation, brain slices were transferred to a recording chamber at room temperature for subsequent recordings.

During the recordings, slices were continuously perfused with ACSF at a flow rate of 3 ml/min. For whole-cell recordings, glass recording pipettes (3–6 MΩ) were prepared using a P-87 glass pipette puller (Sutter Instrument). The pipettes for whole-cell recordings were filled with an internal solution composed of (in mM): 130 K-gluconate, 10 HEPES, 0.6 EGTA, 5 KCl, 3 Na2ATP, 0.3 Na3GTP, 4 MgCl2, and 10 Na2-phosphocreatine, with a pH of 7.2–7.4 and osmolarity of 295–305 mOsm/L.

For neurobiotin microinjection, neurobiotin (2.5 mg/ml; Vector Laboratories, Burlingame, CA, United States) was dissolved in the internal solution. We conducted whole-cell recordings from lateral septum neurons, with the pipette filled with Neurobiotin. After establishing a whole-cell configuration, neurons were held for at least 15 minutes in current clamp and injected with a depolarizing current (500 ms, 500 pA, 1 Hz) to allow for sufficient diffusion of neurobiotin into the cell. At the end of each recording session, the pipette was slowly withdrawn to maintain the cell’s integrity, and the slice was immediately fixed in 4% paraformaldehyde in PBS overnight at 4°C. Post-fixation, slices were washed in PBS and then incubated overnight at 4°C in a solution containing Streptavidin-conjugated Alexa Fluor 488 (1:500 in PBS) to visualize neurobiotin-filled neurons. The slices were then rinsed in PBS, mounted onto slides, and cover-slipped using an antifade mounting medium.

Data were collected using a MultiClamp 700B amplifier (Molecular Devices) and ITC-18 A/D board (HEKA) with AxoGraph software. The neurons were held at −65 mV. The traces were low-pass filtered at 3 kHz and digitized at 10 kHz.The electrophysiological data were analyzed using Easy Electrophysiology. The resting membrane potential was recorded immediately after break-in under I=0 mode. The input resistance of the cell was calculated from the peak of the voltage response to a 200pA hyperpolarized current. The sag or hump amplitude was calculated with a 200pA hyperpolarized current injection. Action potentials were counted using the Action Potential Counting module across current steps from −200 to 200 pA. Action potential properties were analyzed using the Action Potential Kinetics module across current steps from −200 to 200 pA. Threshold was defined as the voltage at the point when the slope first exceeded a value of 20 V/s. Rheobase was defined as the amplitude of the depolarized current injections when the first action potential was observed. Spike half-width is defined as the width at half amplitude. Fast after-hyperpolarization (fAHP) is defined as the difference between the action potential threshold and the minimum voltage after the action potential within 3 ms. Medium after-hyperpolarization (mAHP) is calculated as the difference between the action potential threshold and the minimum voltage after the action potential, from 10 to 50 ms.

### Quantification and Statistical Analysis

Cell counting was done manually using the CellCounter tool in ImageJ. For the lateral septum, the dorsal, intermediate, and ventral regions were designated according to the historical literature (Alonso and Frotscher, 1989). All other regions were designated using the Allen brain atlas. The synaptophysin signal was quantified by measuring the fraction of the total area covered by synaptophysin fluorescence across all brain regions. Basic analysis and plotting were performed using GraphPad’s Prism software or the matplotlib module in Python.

### Microscopy and image analysis

Image acquisition was conducted utilizing a Leica Stellaris 8 confocal microscope, using the 10X, 20X, 40X, and 63X objectives based on the specific requirements of each set of samples. All parameters, such as image acquisition speed, resolution, averaging, zoom, and z-stack, were accordingly adjusted. Post-acquisition, the images were analyzed using the ImageJ software.

## Author contributions

Conceptualization, C.M.R. and C.C.H.; Investigation C.M.R., Y.R., Y.X., M.T.G., D.T, S.V., J.L, M.A.A., S.M.H.; Resources, A.O.P., and C.C.H.; Writing, C.M.R. and C.C.H.; Editing, C.M.R., Y.X., M.T.G., Y.R., C.C.H.; Funding Acquisition, C.C.H.; Supervision, C.C.H.

## Acknowledgements

We thank members of the Harwell lab for feedback on the manuscript; Sandra Chang for lab infrastructure support. We are grateful to the Gord Fishell for providing the N2c-Rabies helper virus and Josh Huang for providing the Lhx2-CreER line. Lastly, we would like to thank the Single Cell Core at Harvard Medical School, Boston, MA for performing the single cell RNA-Seq sample preparation. This research was supported by NIH Grants R37MH119156 and R01NS118933 to CCH and the HHMI Gilliam Fellowship to C.M.R.

---

**Figure S1:**
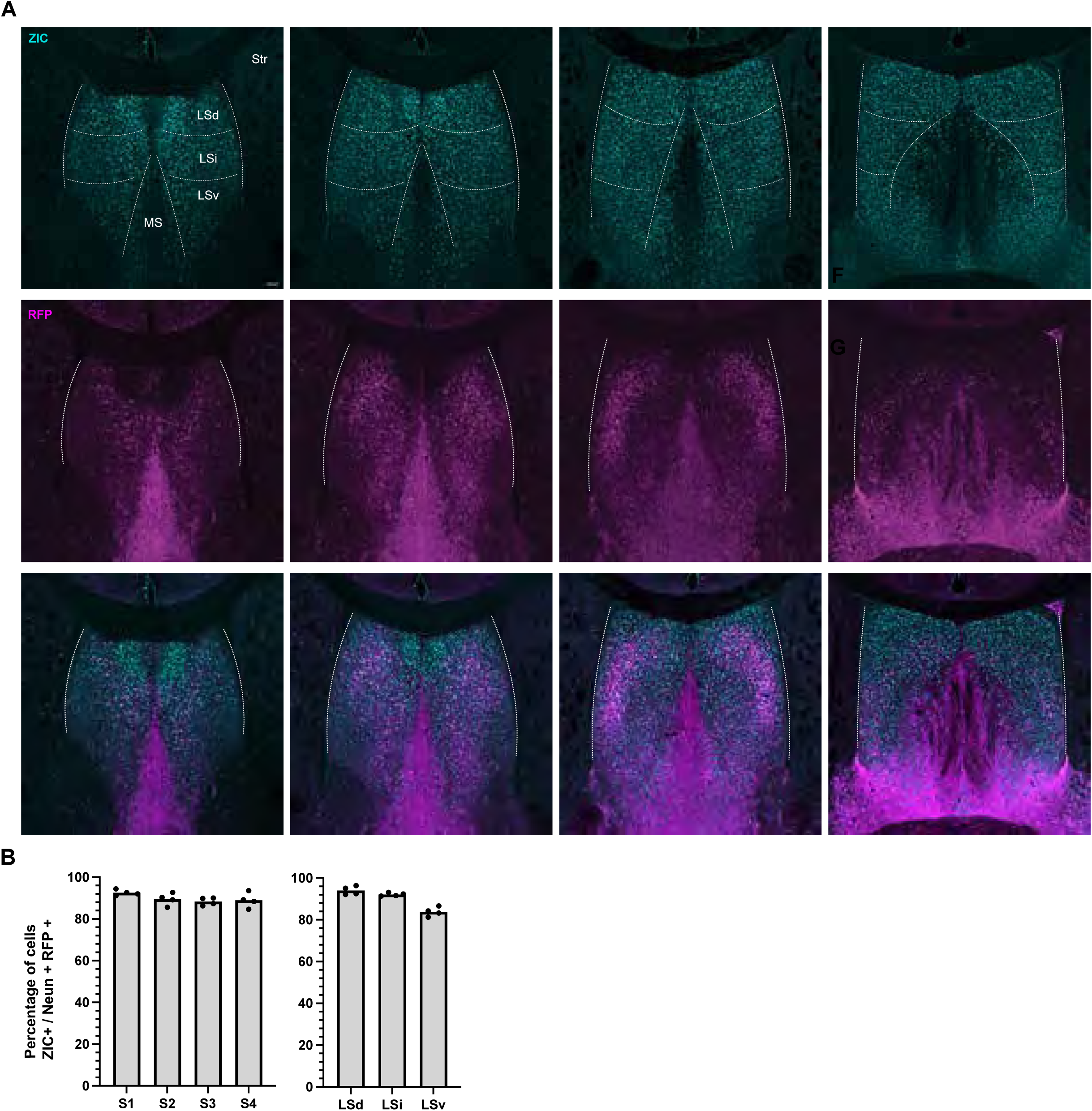
Distribution of *Nkx2.1* lineage cells in the lateral septum. **A.** Coronal sections of the lateral septum across four representative sections along the anterior-posterior axis (AP position from Bregma, from left to right: 0.9, 0.6, 0.3, 0.0) immunofluorescently stained for RFP (magenta) and ZIC (cyan) in a P35 Nkx2.1−Cre animal carrying the Ai14 reporter allele. Scale bar: 100 μm. **B.** Bar graphs showing the proportion of NeuN and RFP positive cells that are ZIC positive, N = 4. The left graph shows counts for the dorsal, intermediate, and ventral subregions of the LS. Right graph shows the counts for the same four regions along the anterior-posterior axis as Figure 1.

**Figure S2:**
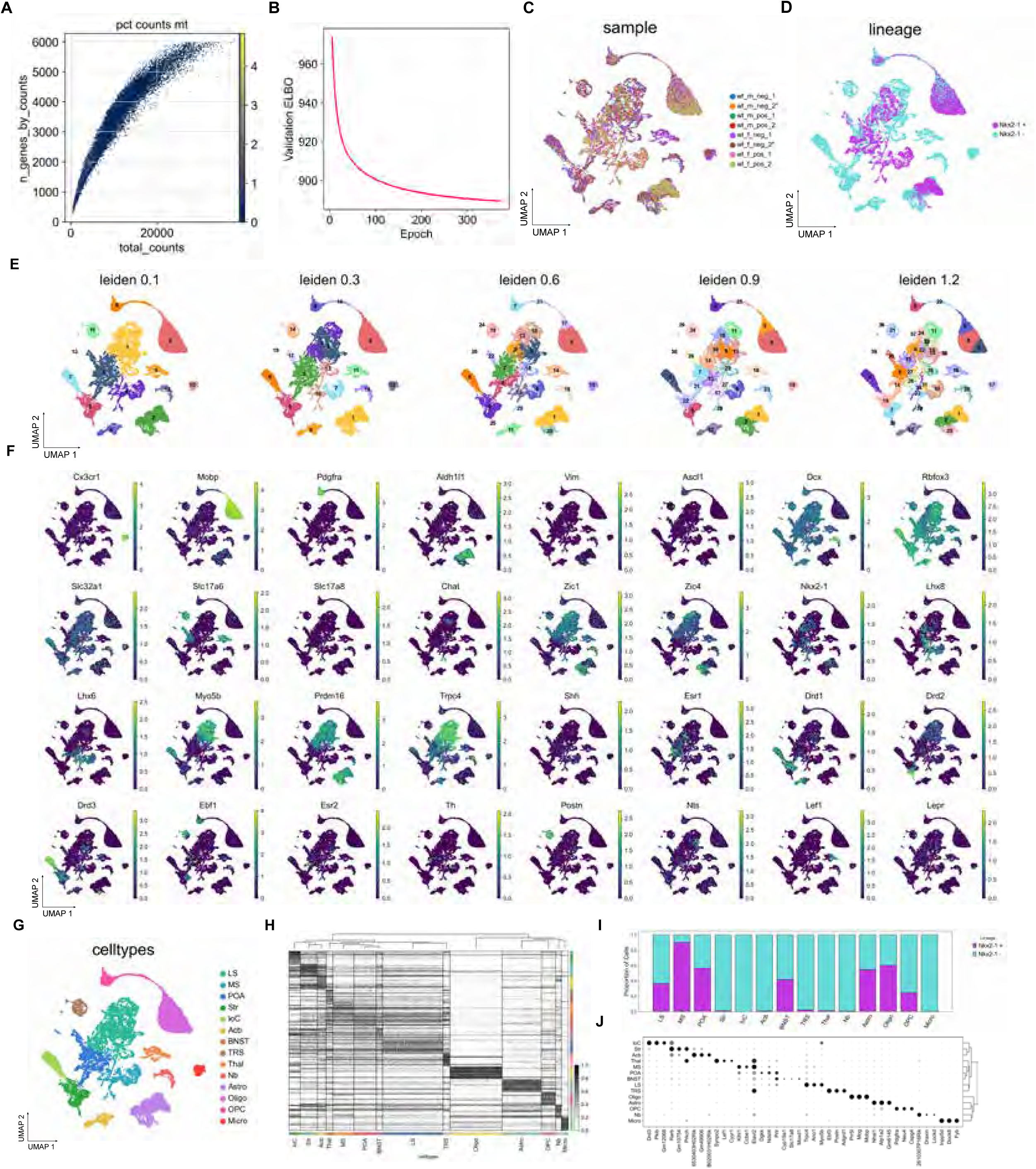
Single-nucleus RNA-seq molecular classification of cell types in the septum. **A.** Scatter plot showing the distribution of raw counts and number of genes for all nuclei. Scale bar indicates the percent mitochondrial genes. **B.** Elbow plot showing the model validation training for the integrated dataset. **C.** UMAP plot showing the balanced mixing of nuclei from different samples. **D.** UMAP plot showing the GFP identity of all 20,678 cells collected (N=6). GFP positive cells (magenta) indicate *Nkx2.1* lineage, while GFP negative cells (cyan) are outside of the *Nkx2.1* lineage. **E.** UMAP plots showing Leiden clusters for cell types at several different resolutions. **F.** Feature plots showing the scaled expression of genes used to identify our major cell groups. **G.** UMAP plot of all cells showing the general cell type annotations. Lateral septum (LS), medial septum (MS), Diagonal band neuron (DbN), Pre-optic area (POA), striatum (Str), Island of Calleja (IoC), Accumbens (Acb), Bed nucleus of the stria terminalis (BNST), triangular septal neurons (TSN), Thalamus (Thal), neuroblasts (Nb), Astrocytes (Astro), oligodendrocytes (Oligo), oligodendrocyte precursor cells (OPC), microglia (Micro).

**Figure S3:**
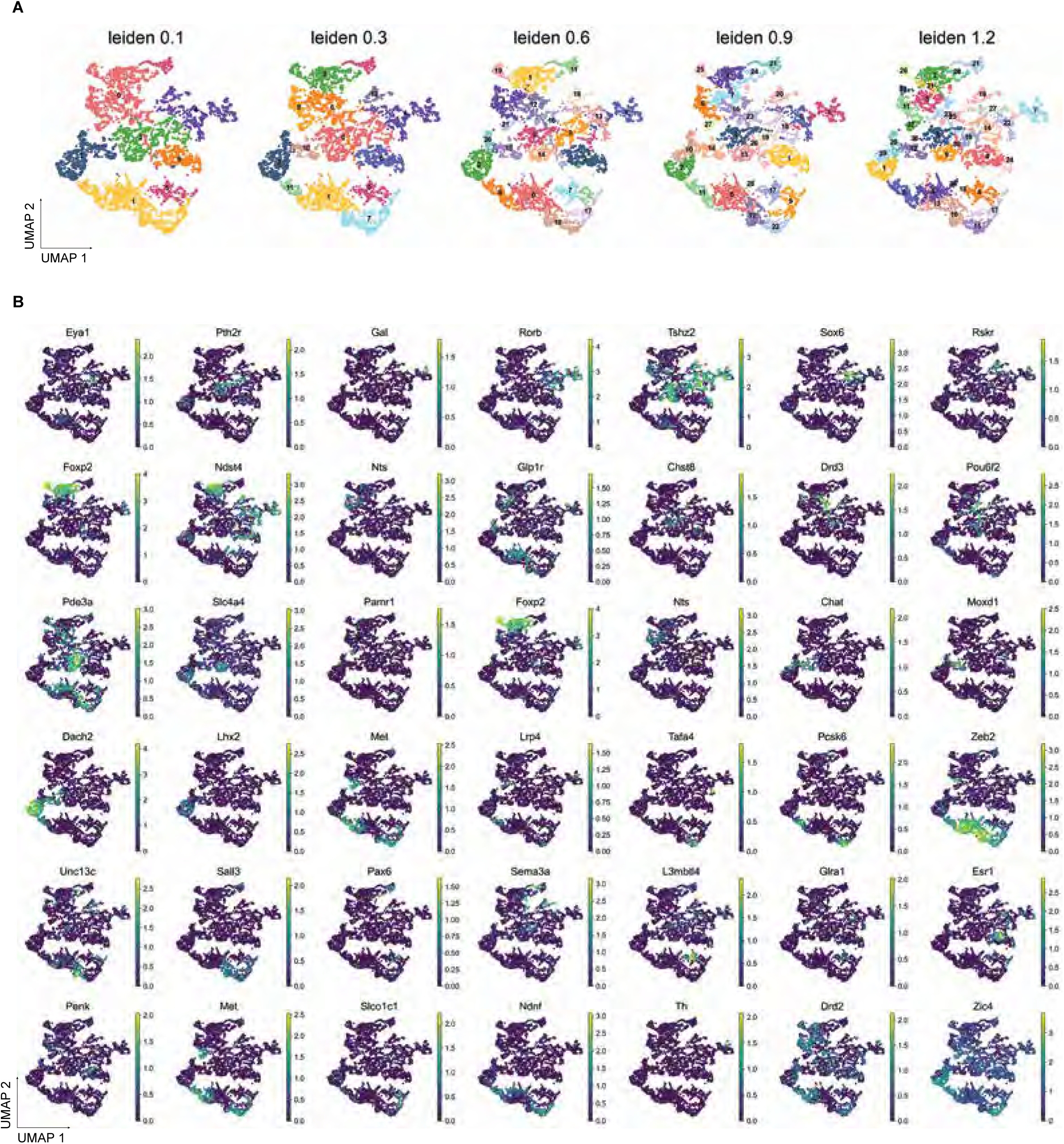
Annotation of lateral septal neuron subtypes. **A.** UMAP plots showing LSNs clustered at several resolutions. **B.** Feature plots of genes that mark LSN subtypes.

**Figure S4:**
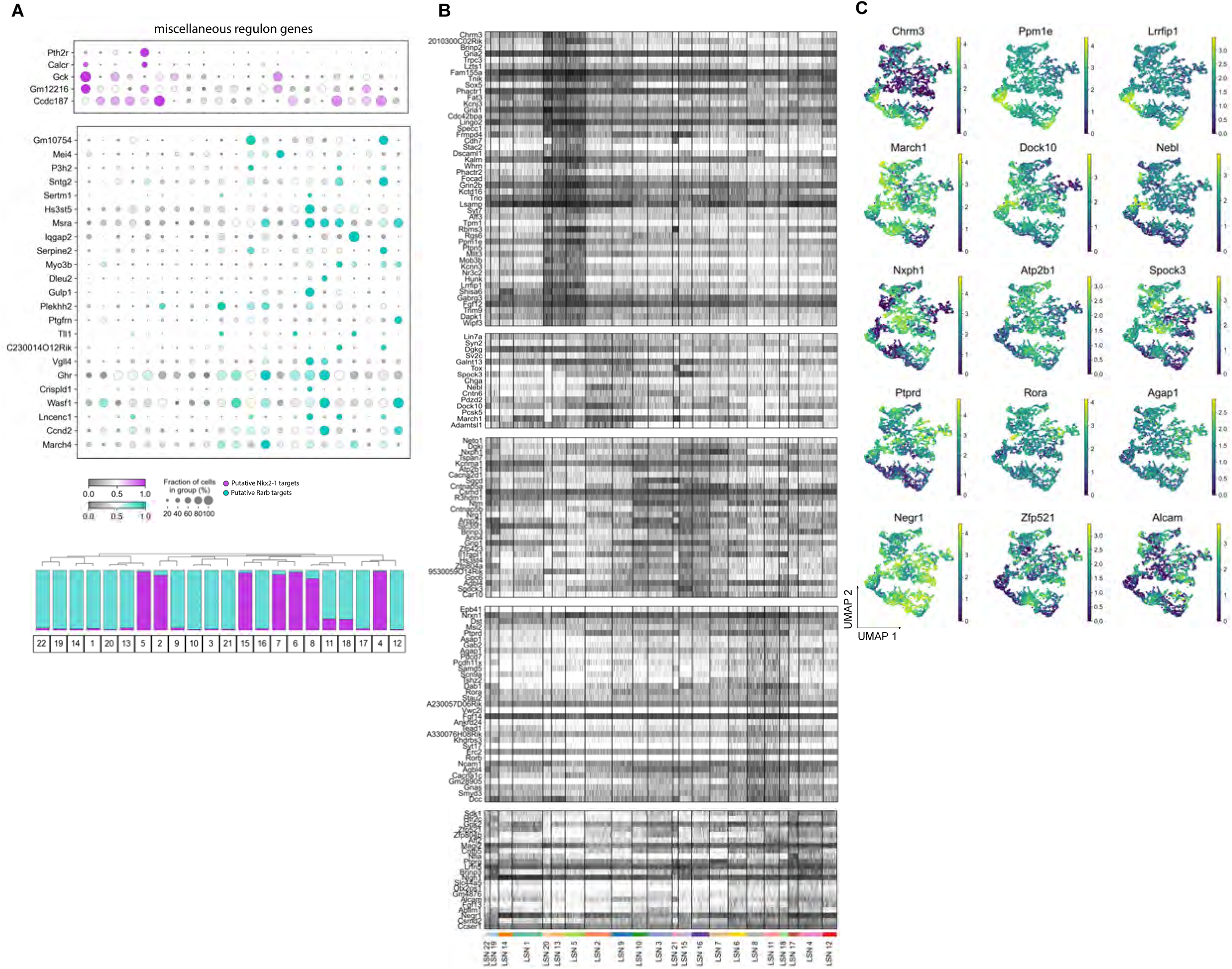
Shared transcriptomic features between developmentally distinct lateral septal neurons. **A.** Dot plot showing the scaled gene expression of miscellaneous genes that shows a lineage bias in expression. **B.** Heatmap showing the top 20 differentially expressed genes for each developmentally distinct paired group. **C.** Feature plot showing scaled gene expression for genes that are shared between developmentally distinct subtypes of neurons.

**Figure S5:**
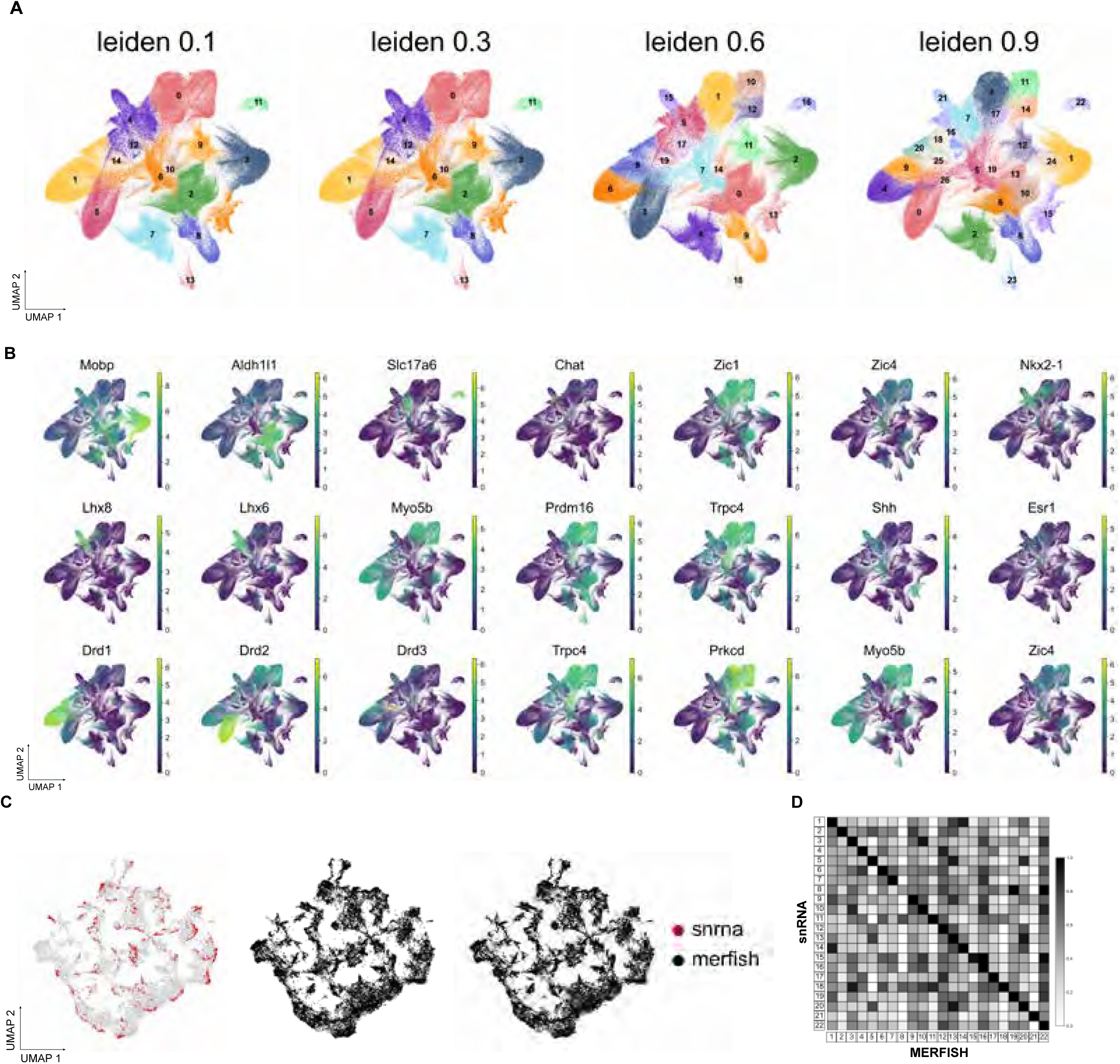
Identification of lateral septal neurons using MERFISH. **A.** UMAP plot showing Leiden clusters at several resolutions for 263,947 cells collected using MERFISH. **B.** Feature plot showing the scaled gene expression of pattern among LSNs for genes specifically enriched in lateral septal neurons. **C.** UMAP plots showing Integration of single-nucleus RNA sequencing data and MERFISH data. **D.** Maxtrix plot showing the correlation between MERFISH and snRNA-seq subtypes.

**Figure S6:**
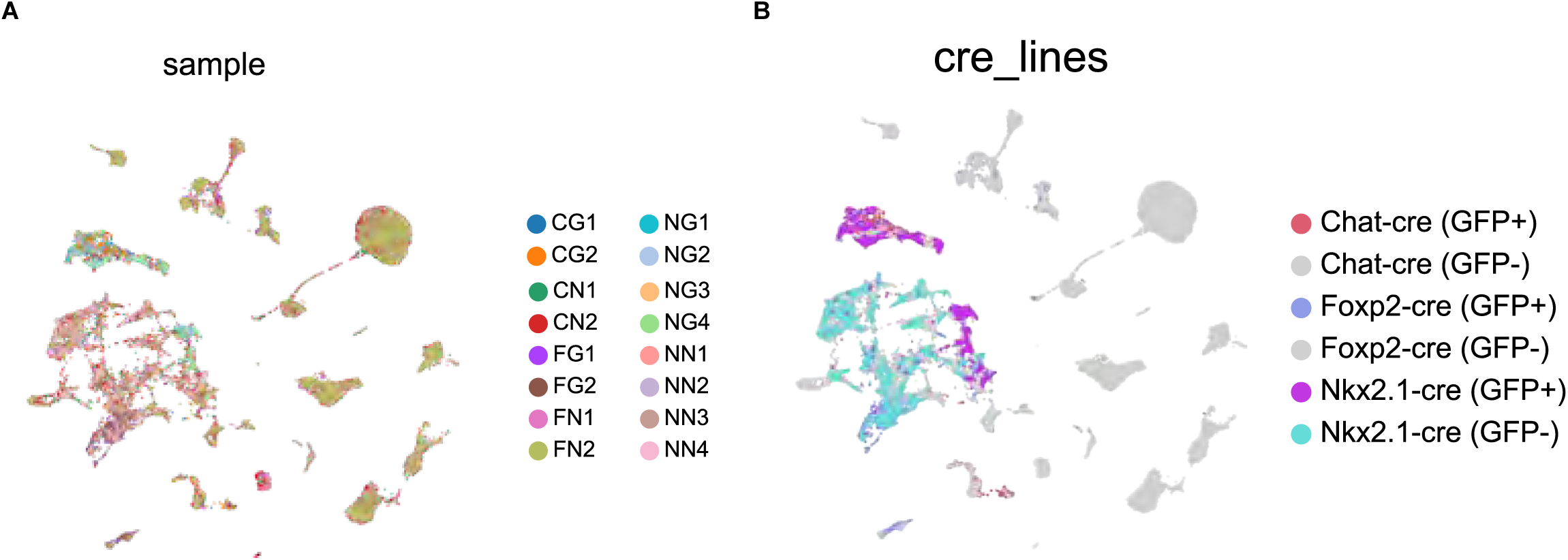
Validation of Cre lines using single-nucleus sequencing. **A.** UMAP plot showing the balanced mixing of nuclei from different samples including samples from the Chat-Cre line and Foxp2-Cre line. **B.** UMAP plot showing cells labeled by each Cre line.

**Figure S7:**
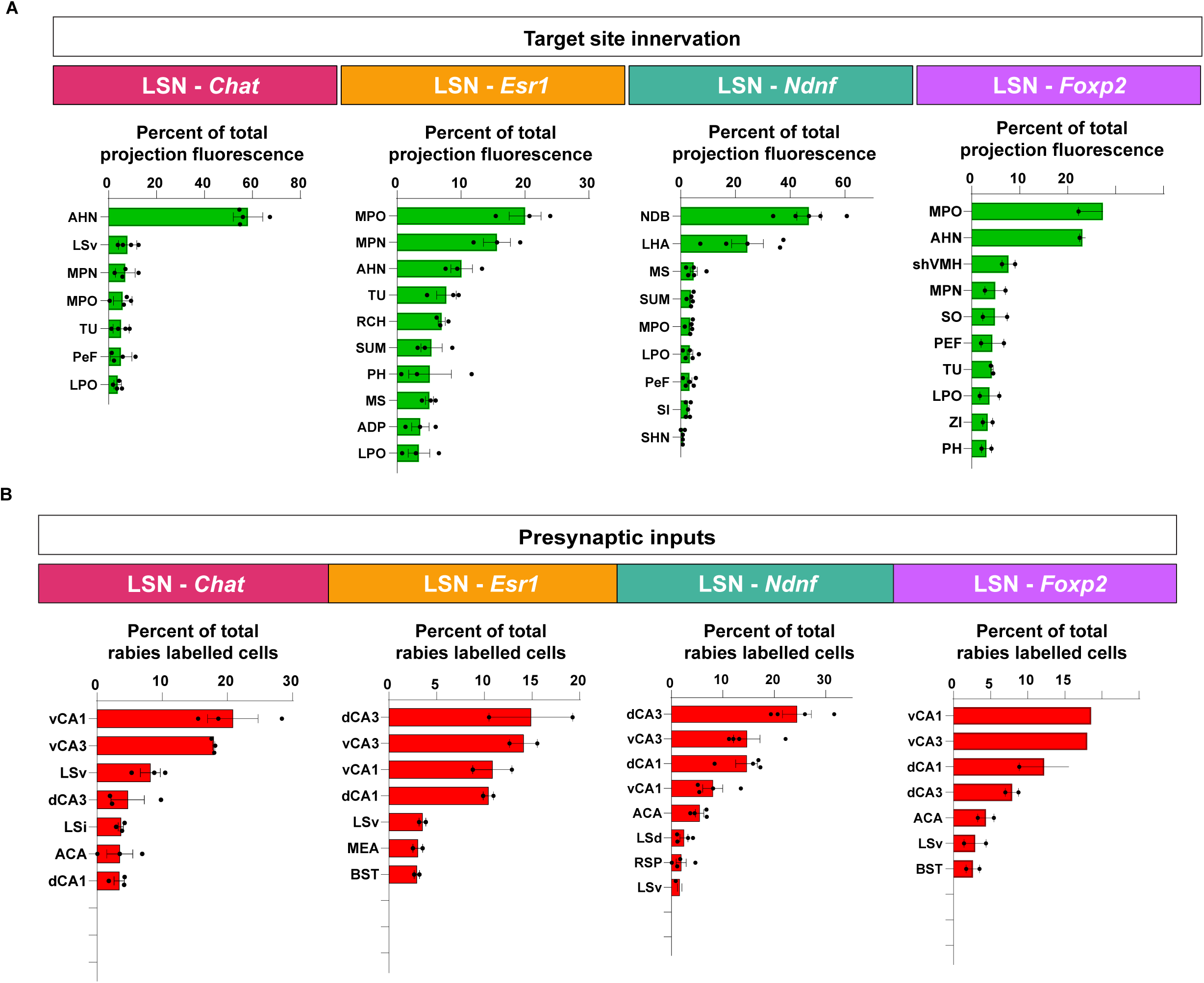
Synaptic inputs and output of lateral septal neuron subtypes. **A.** Quantification of the regional distributions of afferent projections of neurons labelled by different transgenic lines using a local injection of a Cre-dependent AAV. **B.** Quantification of the regional distributions of retrogradely labelled neurons from each transgenic lines.

